# Improving rice drought tolerance through host-mediated microbiome selection

**DOI:** 10.1101/2024.02.03.578672

**Authors:** Alex Styer, Dean Pettinga, Daniel Caddell, Devin Coleman-Derr

**Affiliations:** Department of Plant and Microbial Biology, 111 Koshland Hall, University of California, Berkeley, CA, USA; Plant Gene Expression Center, USDA-ARS, Albany, CA, USA

## Abstract

Plant microbiome engineering remains a significant challenge due to challenges associated with accurately predicting microbiome assembly and function in complex, heterogeneous soil environments. However, host-mediated selection can simplify the process by using plant host phenotype as a reporter of microbiome function; by iteratively selecting microbiomes from hosts with desired phenotypes and using them to inoculate subsequent cohorts of hosts, artificial selection can steer the microbiome towards a composition producing optimized plant phenotypes. In this study, we inoculated rice with wild microbial communities from fallow rice field, desert, and serpentine seep field soils. By challenging these plants with drought and iteratively selecting microbiomes from the least drought stressed plants across multiple generations, we derived simplified microbiomes that enhanced both the growth and drought tolerance of rice. Across selection cycles, microbiomes within and between soil treatments became increasingly similar, implicating both dispersal and selection as drivers of community composition. With amplicon sequencing data we identified specific bacterial taxa associated with improved rice drought phenotypes; while many of these taxa have been previously described as plant growth promoters, we also identified novel taxa exhibiting strong positive correlation with improved drought performance. Lastly, we resolved 272 metagenome-assembled genomes (MAGs) and used these MAGs to identify functions enriched in bacteria driving enhanced drought tolerance. The most significantly enriched functions—particularly glycerol-3-phosphate and iron transport—have been previously implicated as potential mediators of plant-microbe interactions during drought. Altogether, these data demonstrate that host-mediated selection provides an efficient framework for microbiome engineering through the identification of both individual taxa and simplified communities associated with enhanced plant phenotypes.

## Introduction

Plant-associated microbiota directly affect the fitness of their hosts through a variety of mechanisms. These mechanisms include interactions with plant immunity (Pieterse et al. 2014; Durrant and Dong 2004; Schlatter et al. 2017), secretion and metabolism of phytohormones and other signaling molecules (Orozco-Mosqueda, Santoyo, and Glick 2023; Spaepen 2015; Spaepen and Vanderleyden 2011; Glick 2005), mediating nutrient bioavailability (P. Xu and Wang 2023; Richardson and Simpson 2011; Kramer, Özkaya, and Kümmerli 2020), and otherwise altering the physico-chemical environment experienced by plants (Liu et al. 2020; Kumar et al. 2020). Intentional manipulation of plant microbiomes to enhance fitness—through growth promotion, disease suppression, and alleviation of abiotic stress—therefore represents an alternative or supplement to plant breeding to improve agricultural productivity (Friesen et al. 2011; Finkel et al. 2017; Shayanthan, Ordoñez, and Oresnik 2022; Vorholt et al. 2017).

Many individual bacterial and fungal isolates have already been proven capable of plant growth promotion in lab studies (Rodrigues et al. 2008; Bilal et al. 2018; Fan and Smith 2022; Karmakar et al. 2021; Franche, Lindström, and Elmerich 2009), but they often fail to perform similarly well in the field (Sessitsch, Pfaffenbichler, and Mitter 2019). The proximate cause for the performance gap is usually poor colonization and survival (Albareda, Rodríguez-Navarro, and Temprano 2009), but ultimately the discrepancy lies in the difficulty of screening isolates in ways that recapitulate field environments (Russ et al. 2023). In other words, because the strength and direction of a microbe or synthetic community’s (syncom) effect on plant fitness emerge from complex interactions between plants, microbes, and their environment, experiments performed under controlled conditions, at a particular stage of plant development, or in the absence of a native microbial community, fail to capture the spatial and temporal heterogeneity of the field. One way to bridge the gap, at least in part, is to screen for plant growth promotion at the community level—either with wild soil inoculum or a pre-defined syncom—resulting in levels of ecological complexity better approximating field conditions.

Additionally, communities identified as plant-beneficial can include members with synergistic interactions and/or functional redundancy to support both initial colonization and persistence in the soil (Zhuang et al. 2021; Kaur et al. 2022; Liu et al. 2022). Because of these synergies and functional redundancy, syncoms may even have greater resilience to functional shifts across environmental heterogeneity compared to single-isolate inoculations alone (Bradáčová et al. 2019).

However, syncoms are often designed around *a priori* knowledge of individual isolates rather than taking a community-oriented approach (Vorholt et al. 2017). Isolate-first approaches usually aim to saturate a syncom with individuals known to perform a specific probiotic function, and might additionally screen additional isolates for stable pairwise interactions to help support probiotic engraftment (Herrera Paredes et al. 2018; Faust and Raes 2012; Friedman, Higgins, and Gore 2017). Nearing a community-level approach, others have attempted to traverse the isolate sample space by recapitulating simplified versions of naturally assembled communities (B. Niu et al. 2017), or otherwise formulating syncoms around known patterns of co-occurrence or syntrophy (H. Wang et al. 2017; Jones and Carlson 2019; McCarty and Ledesma-Amaro 2019). While these approaches have resulted in syncoms with demonstrated plant-beneficial effects (e.g. Herrera Paredes et al. 2018), they quickly become prohibitively time and labor-intensive as the number of isolates to be screened increases.

We propose that host-mediated selection has been an underutilized framework as a starting point for microbiome engineering, one that inherently accounts for ecological interactions within a microbiome without requiring *a priori* knowledge about them. In brief, host-mediated selection uses host phenotype as a reporter to artificially select microbiomes associated with desirable host phenotypes (U. G. Mueller and Sachs 2015; Ulrich G. Mueller and Linksvayer 2022). By targeting the emergent property of a microbiome (i.e. its impact on host phenotype), selection occurs at both the individual microbe and microbial community levels to maintain or exclude specific taxa and interactions (Whitham et al. 2020). In other words, host-mediated selection not only identifies taxa with direct benefit to host phenotype, but also identifies taxa that indirectly benefit their plant hosts via their interactions with other microbiome members.

Here, we describe a long-term experiment where we applied host-mediated selection to derive simplified root microbiomes that enhance rice drought tolerance. Drought severely reduces rice production and threatens the food security of billions of people around the globe (Panda, Mishra, and Behera 2021; Pandey and Shukla 2015). On the other hand, cultivation in flooded paddies emits significant amounts of methane and nitrous oxide, leading rice to account for nearly half of greenhouse gas emissions from croplands (Qian et al. 2023). Consequently, discovering new ways to cultivate rice with less water can be of benefit to both people and the environment. In addition to identifying bacterial communities and individual taxa with the potential to enhance rice drought tolerance, we also contribute to the nascent body of host-mediated selection (HMS) literature by tracking the HMS process with large amounts of sequence data (>1,100 samples with 16S rRNA sequence data, combined with WGS data) and being the first of its kind to compare outcomes of the same HMS process applied to diverse sources of inocula in parallel.

## Results

To start, we established four soil treatments, three of which were inoculated with diverse field-collected soil microbiota; the fourth treatment received no initial inoculation and was left to acquire microbes from the environment alone. Each soil treatment further consisted of multiple selection and control lines (**Figure 1**). Each Selection Line (SL) included 15 plants per generation from which 3 were selected and their microbiomes harvested to inoculate a subsequent generation of 15 plants. Importantly, selection lines were independent of one another, and “generation” refers to a 40 day selection cycle rather than a seed to seed cycle. No drought treatment was applied in the first generation, with the intent to simply enrich microbiota associated with healthy rice roots to start. Accordingly, we refer to this selection cycle as the Enrichment Generation (EG) and subsequent selection cycles as Selection Generations (SG), during which rice plants were challenged with a 10 day drought treatment after 30 days of well-watered growth.

**Figure 1.**
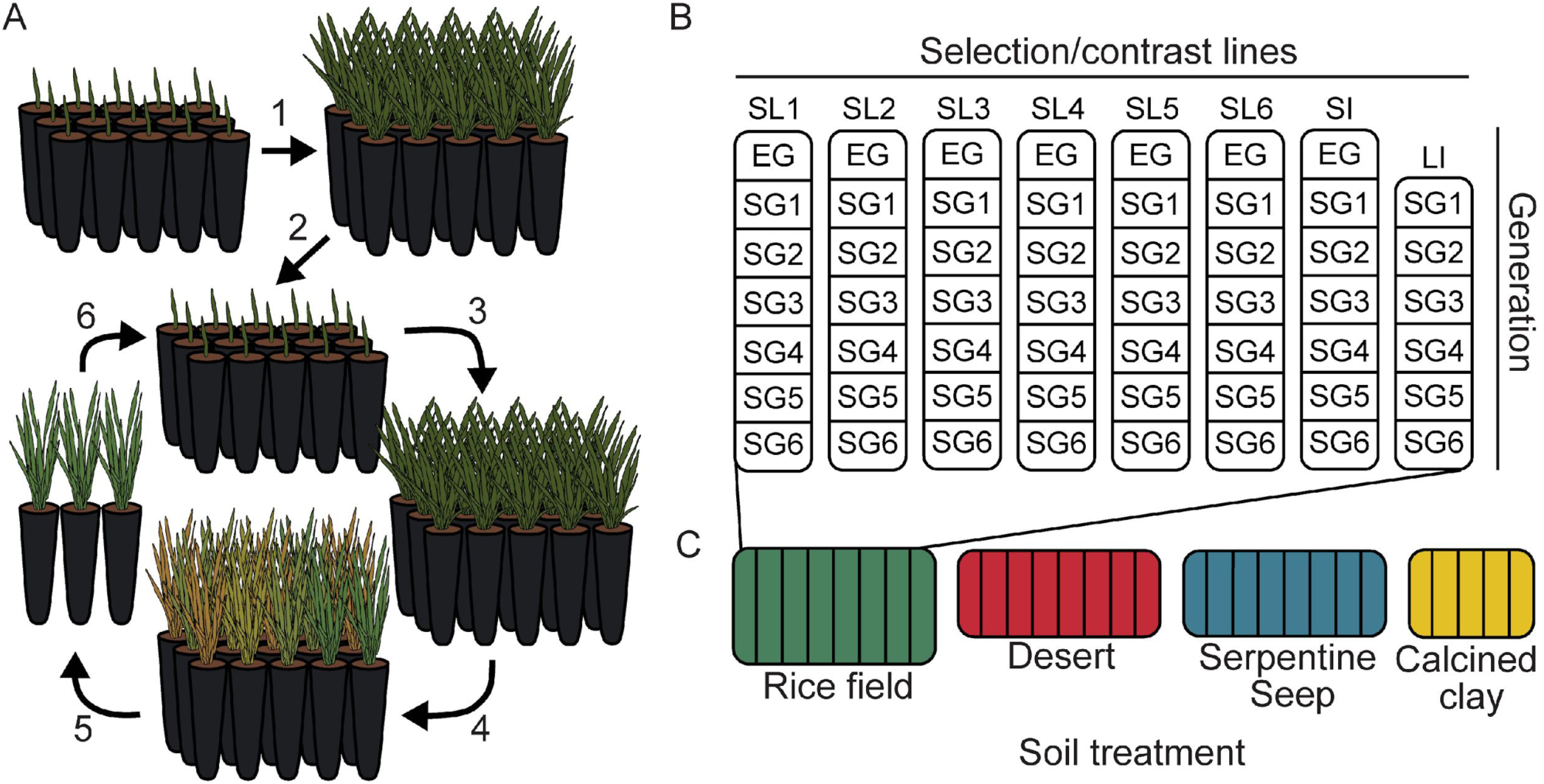
Host-mediated selection experimental design. (**A**) Representative of a single selection line of 15 rice plants. 1) Enrichment Generation (EG): rice are transplanted into 1:1 mixtures of field soil and sterilized calcined clay grow for 40 days in well-watered conditions. 2) Roots and adhering soil are harvested, minced, and ground, then mixed into sterilized calcined clay to inoculate the first selection generation (SG1). 3) SG plants grow for 30 days in well-watered conditions. 4) Drought treatment is initiated; plants are imaged daily to track phenotypes for 10 days. 5) The three best-performing plants are identified by drought scores derived via Normalized Difference Vegetative Index (NDVI) and biomass data. 6) Roots and soils from best-performing plants are harvested and diluted with sterilized calcined clay to inoculate the subsequent selection generation (SG2). Steps 3-6 are then repeated iteratively for additional selection generations. (**B**) An overview of the type and number of selection and control lines for an individual soil treatment. Selection Lines (SL 1-6) consist of 15 plants per generation each and were maintained as independent lines across generations (i.e. transfers of selected inocula occurred within SLs). Sterile- and Live-Inoculated control lines (SI and LI), on the other hand, each included 30 plants per generation and were inoculated by an amalgam of material from all selection lines within their soil treatment; SI inocula were autoclave-sterilized prior to each generation. Like their SL counterparts, SI plants were well-watered throughout the enrichment generation and subjected to drought in selection generations; LI plants were well-watered in all generations. (**C**) Rice Field (green), Desert (red), and Serpentine Seep (blue) soil treatments—inoculated by field soils—each included six selection lines (SL 1-6) of 15 plants per generation. Calcined Clay (yellow) did not receive any inoculum in EG, instead assembling microbiomes composed of environmental taxa, and included just three SLs of 15 plants per generation. The number and length of columns are proportional to the number of lines and selection generations, respectively, associated with each soil treatment.

Each soil treatment also included two control lines, one Sterile-Inoculated (SI) and the other Live-Inoculated (LI). In contrast to selection lines, each of the two control lines included 30 plants per generation, and microbiomes were not selected for propagation forward into future generations. Instead, both control lines were intended to provide meaningful contrasts to selection lines: SI underwent the same drought treatment as selection lines, but received sterilized versions of selection line inocula; LI were never droughted but received unsterilized inocula from selection lines. Simply put, SI plants were meant to serve as a static control for rice drought performance across generations; LI plants were meant to help identify microbes enriched by drought specifically, and to determine whether microbiomes optimized for drought were deleterious to the performance of unstressed plants (See **Figure 1** and Methods for greater detail on experimental design, rice phenotyping methods, inoculum preparation, and selection criteria). Comprehensive reviews of host-mediated selection experimental design and terminology can also be found in Mueller and Sachs 2015 and Mueller and Linksvayer 2022 (U. G. Mueller and Sachs 2015; Ulrich G. Mueller and Linksvayer 2022).

### Source inocula bacterial diversity

Without specific *a priori* expectations of which microbial taxa or functions would be required to improve drought tolerance in rice, we first screened a panel of nine field soils from distinct ecoregions and land-use histories across California for rice growth phenotypes. Each soil was characterized by unique physico-chemistry and bacterial diversity, even at broad taxonomic levels (**Figure 2, Supplemental Figure 1, Supplemental Table 1**). And, after inoculating rice plants with each of the soils, we observed differences in rice growth phenotypes between soil inoculation groups (**Supplemental Figure 1D-F**). Of the field soils whose microbes better supported rice growth, we chose three to use as source inocula for host mediated selection: Rice Field, from a fallow rice field; Desert, from the Mojave Desert; and Serpentine Seep, from a serpentine soil that experiences consistent groundwater seep. In addition to their relatively beneficial impact on rice plants within our pilot experiment, we chose these soils for their negligible overlap in taxonomic composition, which would provide host-mediated selection with three distinct starting points. Further, we reasoned that each soil community might have its own set of advantages: Rice Field microbes might be more likely to form associations with rice roots and tolerate flooded conditions; Desert taxa might be better adapted to drought; and the high diversity of Serpentine Seep taxa, in terms of both richness and evenness, might provide more functional diversity for host-mediated selection to act upon. Just prior to host-mediated selection, we collected fresh field samples of Rice Field, Desert, and Serpentine Seep soils. For good measure, we characterized the bacterial diversity of these new collections, the results of which were consistent with what we had previously observed (**Figure 2C, Supplemental Figure 1**). The fourth soil treatment included in host-mediated selection, Calcined Clay, received no field soil inoculum; as the name of this treatment suggests, the first generation of these plants was sown directly into sterilized calcined clay substrate.

**Figure 2.**
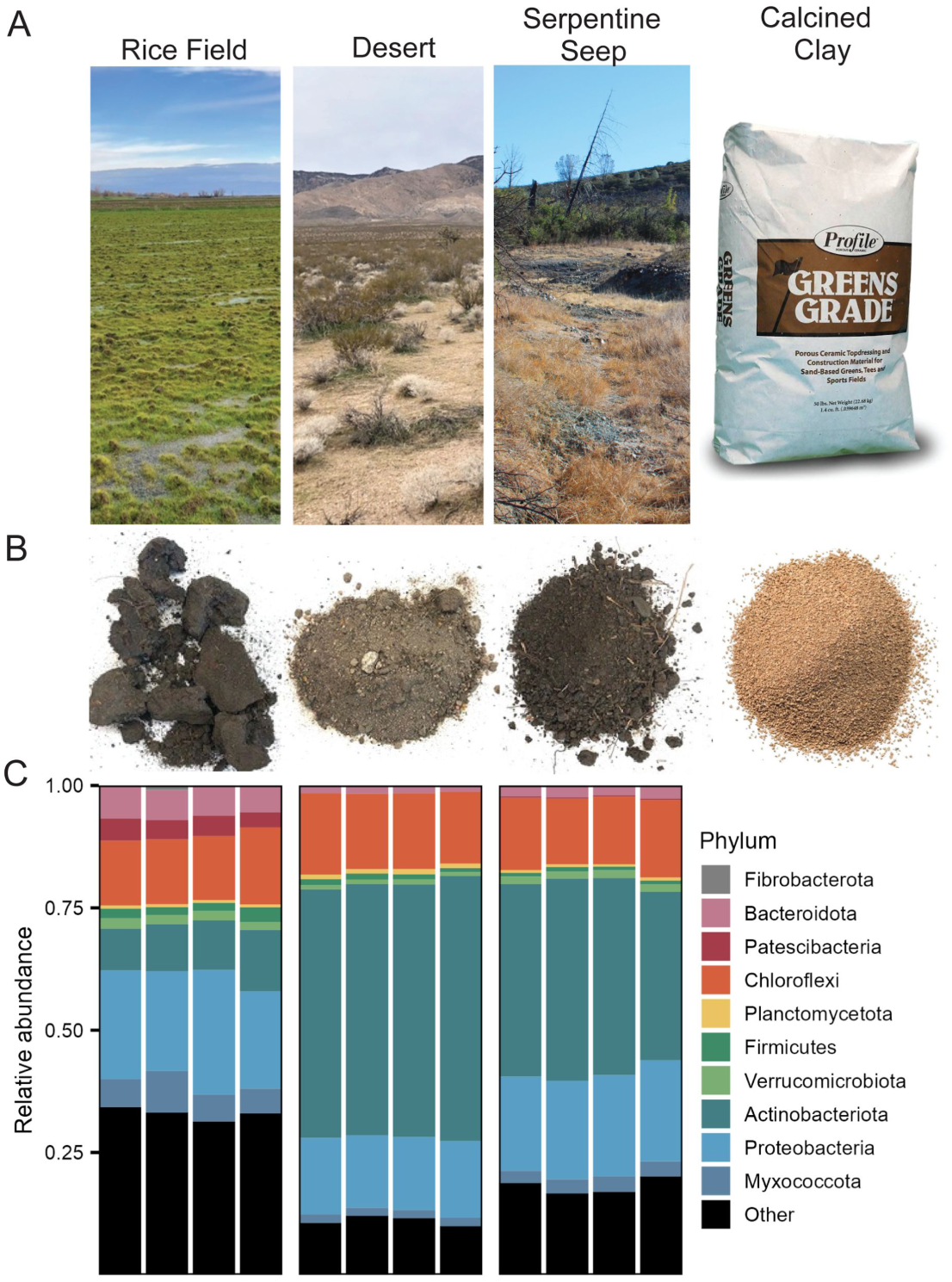
Field sites, soils, and starting diversity of source inocula. **(A)** Collection sites for Rice Field, Desert, and Serpentine Seep. Calcined Clay “source inocula” was simply calcined clay (Profile Greens Grade, Cat.No. FZ-TP) that was rinsed and autoclave-sterilized before use. **(B)** Field soils differed in physico-chemical characteristics (Supplemental Table 1). Rice Field was visibly clay-dominant, Desert predominantly sandy, and Serpentine Seep high in organic matter, but less clay-like compared to Rice Field. **(C)** Bacterial diversity and community compositions were distinct for each soil, even at broad taxonomic levels (Supplemental Figure 1).

### Rice phenotypes improve across selection generations

To score drought tolerance in our system, we created a non-destructive, image-based protocol to measure the Normalized Difference Vegetation Index (NDVI) of individual plants during drought. Briefly, NDVI is a metric that quantifies plant stress by measuring the intensities of red and infrared light reflected by leaf surfaces and is most commonly used in remote sensing to determine plant health (Rouse et al. 1974; Huang et al. 2021). Prior to host-mediated selection we performed an additional pilot experiment confirming that measured values of NDVI drop as drought intensity increases (**Supplemental Figure 2D**); these values also had strong correlation with plant shoot percent water content (*R*=0.98, *p*<2.2e-16; **Supplemental Figure 2E**). To derive a final drought score, we calculated the area under the curve of NDVI throughout the drought treatment and adjusted these scores to remove the effect of plant biomass.

Additional information about our protocol can be found in the Methods as well as in

**Supplemental Figure 2**.

In the context of host-mediated selection, we observed consistent, significant increases in drought score across selection generations for Rice Field, Desert, and Calcined Clay soil treatments, but not for Serpentine Seep (**Figure 3**). Specifically, drought scores significantly increased in all selection lines (SLs) belonging to Rice Field (6/6), Desert (6/6), and Calcined Clay (3/3), but in just two SLs belonging to Serpentine Seep (2/6) (**Figure 3B, Supplemental Figure 3A**). In further contrast to the improved performance of other soil treatments, two Serpentine Seep SLs saw significantly lower drought scores across selection generations (**Supplemental Figure 3A**).

**Figure 3.**
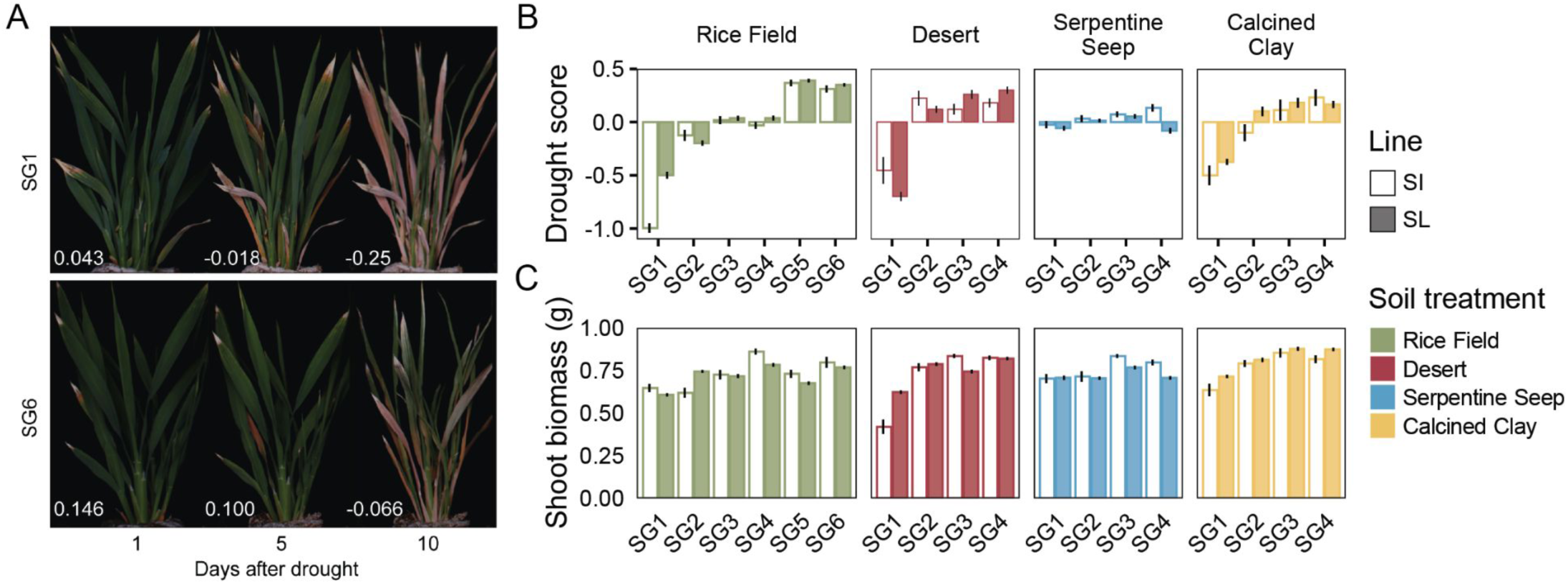
Rice phenotypes across host-mediated selection generations. (**A**) Representative images of Rice Field selection line plants in the first (SG1) and last (SG6) selection generations at 1, 5, and 10 days after drought; drought scores are shown in the bottom left corner of each image. (**B**) Drought score, quantified as the area under the curve of NDVI values, significantly improved for all soil treatments except Serpentine Seep. Here, we show data for all selection lines combined (gray bars) and assessed the response of drought score to generation via linear mixed effects models (drought score ∼ generation + (1|selection line); Rice Field: β=0.17, *t* = 28.8, *p* < 2e-16; Desert: β=0.19, *t* = 14.1, *p* < 2e-16; Serpentine Seep: β=-0.003, *t* = -0.3, *p* = NS; Calcined Clay: β=0.17, *t* = 8.7, *p* < 0.001). Filled bars represent the mean drought score of all selection line (SL) plants +/- one standard error; white, unfilled bars are data for sterile-inoculated (SI) plants. Because selection lines were independent of one another, we also tested them individually via OLS regression: all selection lines of Rice Field (6/6), Desert (6/6), and Calcined Clay (3/3) significantly improved over time; Serpentine Seep included selection lines that both significantly improved (2/6) and worsened (2/6) (Supplemental Figure 3). (**C**) Shoot dry weight biomass also significantly increased across selection generations for all soil treatments, except Serpentine Seep (Rice Field: β=0.02, *t* =7.0, *p* < 2e-16; Desert: β=0.06, *t* =11.6, *p* < 2e-16; Serpentine Seep: β=0.006, *t* =1.2, *p* = NS; Calcined Clay: β=0.05, *t* =10.3, *p* < 2e-16). Tested individually, all selection lines of Rice Field (6/6) and Calcined Clay (3/3) significantly improved over time; most Desert selection lines (5/6) significantly improved; Serpentine Seep included one selection line that significantly increased (1/6) and one that significantly decreased (1/6) (Supplemental Figure 3).

Notably, the improvements in drought score of non-Serpentine Seep soil treatments did not come at the expense of plant biomass (**Figure 3C**). Rather, shoot biomass generally increased over time: all Rice Field (6/6) and Calcined Clay (3/3) SLs, and most Desert SLs (5/6), significantly increased shoot biomass across selection generations. Two Serpentine Seep SLs showed significant change over time, with one (1/6) increasing and the other (1/6) decreasing in biomass (**Supplemental Figure 3B**). Sterile-inoculated (SI) control lines of all soil treatments, Serpentine Seep included, also saw significant increases in both drought score and biomass phenotypes over time (**Figure 3, Supplemental Figure 3**). Though each SI plant received sterilized inoculum and was maintained as a spatially discrete unit (**Supplemental Figure 4**), they were not axenic; consequently, SI plants were colonized by environmental microbiota throughout each 40-day generation. Overall, given that environmental conditions and rice genotypes were held constant across selection generations, these data suggest that changes in microbiome composition and function were driving improvements in rice drought phenotypes, even in SI plants.

### Microbiome diversity and composition over time

To assess the impact of host-mediated selection on bacterial community composition and diversity across selection generations, we performed 16S rRNA sequencing on all root samples from Rice Field (*n* = 1019) as well as a subset of those from Desert (*n* = 58), Serpentine Seep (*n* = 58), and Calcined Clay (*n* = 42) soil treatments. At the start, high microbial diversity was observed across all soils, Serpentine Seep exceptionally so, with higher levels of species richness (2507 observed ASVs) and evenness (Shannon Index = 7.1) compared to Rice Field (1390 observed ASVs; Shannon Index = 6.3) and Desert soils (869 observed ASVs, Shannon Index = 6.1) (**Figure 4A-B**). Within several selection generations, however, bacterial communities of all soil treatments had been reduced to just ∼200 ASVs. While diversity decreased steadily over time in Rice Field microbiomes, both Desert and Serpentine Seep lost most of their input diversity in the enrichment generation (EG) alone (**Figure 4A-B**). Considering that only roots from EG plants were harvested to inoculate plants in SG1, the large reduction in diversity during EG is consistent with the observation that microbial diversity generally declines along transects spanning from bulk soil to root endophytic spaces (Kuzyakov and Razavi 2019). Additionally, Rice Field better maintained diversity in this first generation, supporting our initial hypothesis that this soil would be more likely to include taxa that associate with rice roots. In contrast to the field soil treatments, Calcined Clay SL microbiomes saw modest increases in diversity between EG (78 observed ASVs; Shannon Index = 2.9) and SG4 (138 observed ASVs; Shannon Index = 3.6), consistent with the fact that this treatment did not receive a field soil inoculum in EG, but was allowed to recruit microbiota from the growth chamber environment (**Figure 4A-B**). Similarly, sterile-inoculated plants successfully recruited microbes from the environment each generation, though the resulting communities were consistently less diverse compared to their selection line counterparts (**Figure 4A-B**).

**Figure 4.**
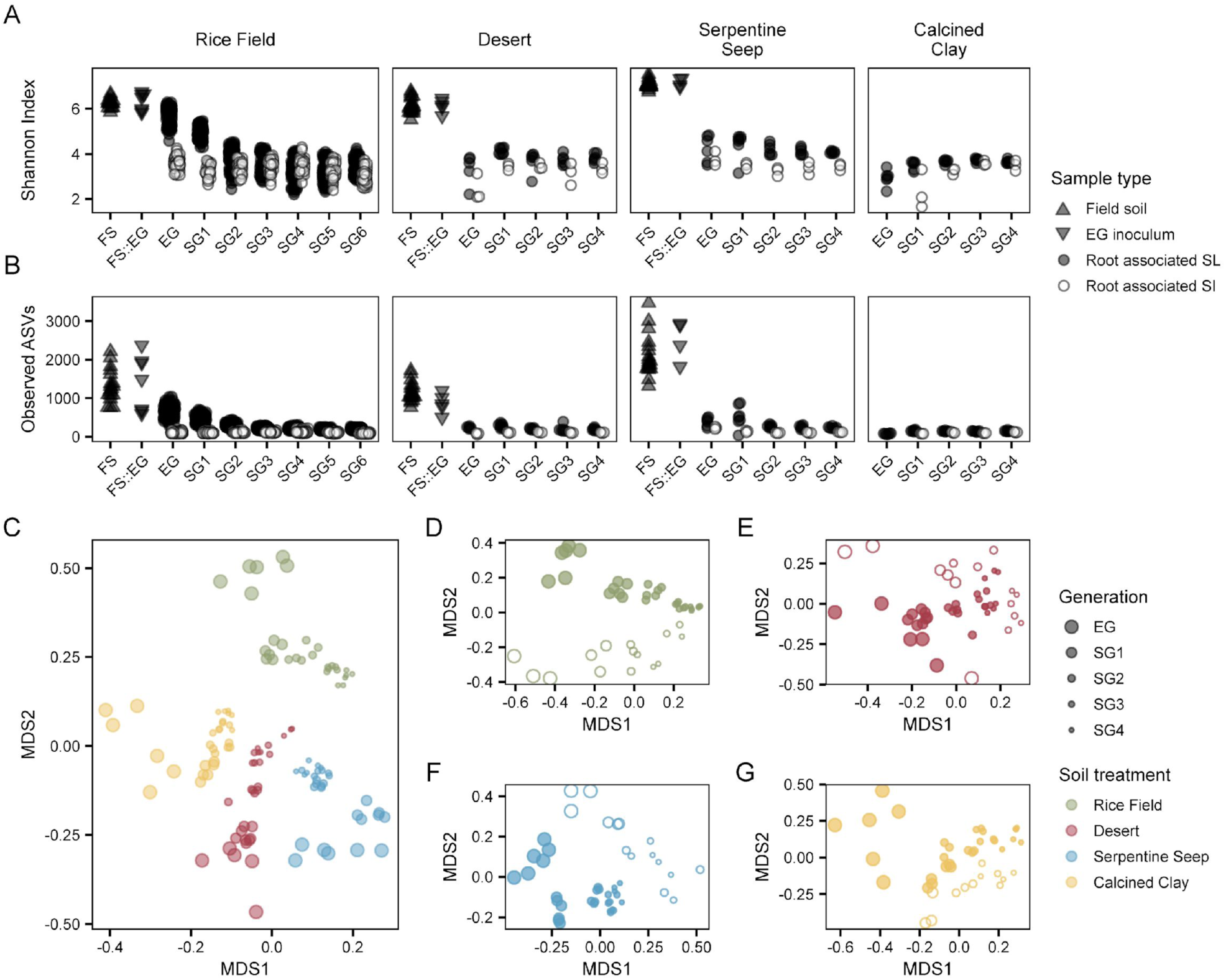
Alpha and beta diversity of bacterial communities over time. (**A**) Shannon-Index diversity and (**B**) number of observed ASVs for soil treatments over time. Each point represents a bacterial community from one of several sample types, including original field soils (FS), enrichment generation (EG) inocula (FS::EG), root-associated selection line (SL), or root associated sterile-inoculated (SI); sample types are indicated by shape. EG inocula were simply field soil diluted by sterilized calcined clay. Sample sizes and numbers of selection generations differ between soil treatments; having observed increasing similarities in microbiome composition between soil treatments after four selection generations, we decided to focus effort and resources on Rice Field, which we carried forward an additional two selection generations. (**C**) Bray-Curtis (BC) dissimilarity between soil treatment selection lines plotted in two-dimensional space. Each point represents an individual sample, where color indicates soil treatment and size indicates generation. BC values are the average of 1000 permutations where each sample was individually subset to 5000 reads. (**D-G**) Similar to (C), but panels are individual soil treatments and include both SL (filled) and SI (unfilled) samples. Despite forming distinct clusters, SL and SI microbiomes generally follow parallel compositional trajectories through time. For ordinations in panels C and D, Rice Field samples were subset to match the sequencing effort of other soil treatments prior to calculating BC values.

Next, we assessed the impact of host-mediated selection on bacterial community beta diversity using Bray-Curtis dissimilarity, which is particularly useful for showing changes in community composition along a gradient or through time (Anderson et al. 2011). Though soil treatments maintained distinctive microbiome identities throughout the experiment, their community compositions became increasingly similar across selection generations (**Figure 4C, Supplemental Figure 5A,C**). Within each soil treatment, SL and SI bacterial communities formed distinct clusters, but changes in their composition also followed similar trajectories over time (**Figure 4D-G**). Because Bray-Curtis is not a true distance metric and required rarefaction to normalize read counts between samples, we additionally quantified Aitchison distance which does not rely on rarefaction and is explicitly intended for compositional data (Martino et al. 2019). Both metrics were generally in agreement and confirmed that microbiomes of different soil treatments increased in similarity over time (**Supplemental Figure 5A,C**). Within field soil treatments—Rice Field, Desert, and Serpentine Seep—both metrics also show SL microbiomes becoming more similar to their sterile-inoculated counterparts over time, with the exception of Desert which is static over time by Aitchison distance (**Supplemental Figure 5B,D**). Differences between Calcined Clay SL and SI, on the other hand, were static when measured by

Bray-Curtis, and increased over time when measured by Aitchison distance. To confirm that the overall increase in similarity between soil treatments was not solely driven by shared sparsity, we simply counted the number of ASVs shared between SL plants of each soil treatment: in the enrichment generation, just 0.4% (28/7106) of observed ASVs were shared by three or more soil treatments, increasing to 4.3% (107/2471) by SG4 (**Supplemental Figure 6A**). Moreover, shared ASVs tended to be more abundant, with a subset of just 20 shared ASVs eventually accounting for up to 60% of microbiome composition in SG4 (**Supplemental Figure 6B**).

### Dispersal and growth rate influence bacterial fitness

Given the trend towards more uniform community composition both within and between soil treatments over time, we sought to better understand whether shared ASVs represented a common set of taxa recruited from the growth chamber environment or if taxa dispersed between soil treatments. To do so, we assigned ASVs as originally belonging to one or multiple soil treatments based on which original field soils and EG inocula they were observed in.

Mapping these designations onto the bacterial community data, it became clear that shared ASVs were the result of dispersal events between soil treatments as well as recruited from the environment (**Figure 5**). Environmental bacteria quickly rose to high abundance in all soil treatments—as much as 50% of relative abundance by the end of SG1—and remained a significant fraction of the community throughout subsequent selection generations. Despite the colonization of non-native bacteria, the native diversity of Rice Field and Serpentine Seep SL microbiomes remained the dominant fraction, whereas the native diversity of Desert SL microbiomes steadily declined. Calcined Clay microbiomes, initially uninoculated, were primarily composed of environmental taxa with the remaining fraction evenly represented by treatments inoculated with field soils. Interestingly, microbiomes of SI plants were biased towards the diversity of their respective soil treatments, though the fraction of native diversity in these communities was consistently less compared to their SL counterparts (**Figure 5**); this bias was likely the result of dispersal limitation, priority effects, or both, given that plants were regularly shuffled within—but not between—soil treatments during each generation. Overall dispersal proved to be a fitness advantage in the context of host-mediated selection by increasing a taxon’s likelihood of being observed in a selected microbiome and allowing re-inoculation in lines where the taxon was lost due to either host-mediated selection or stochastic drift.

**Figure 5.**
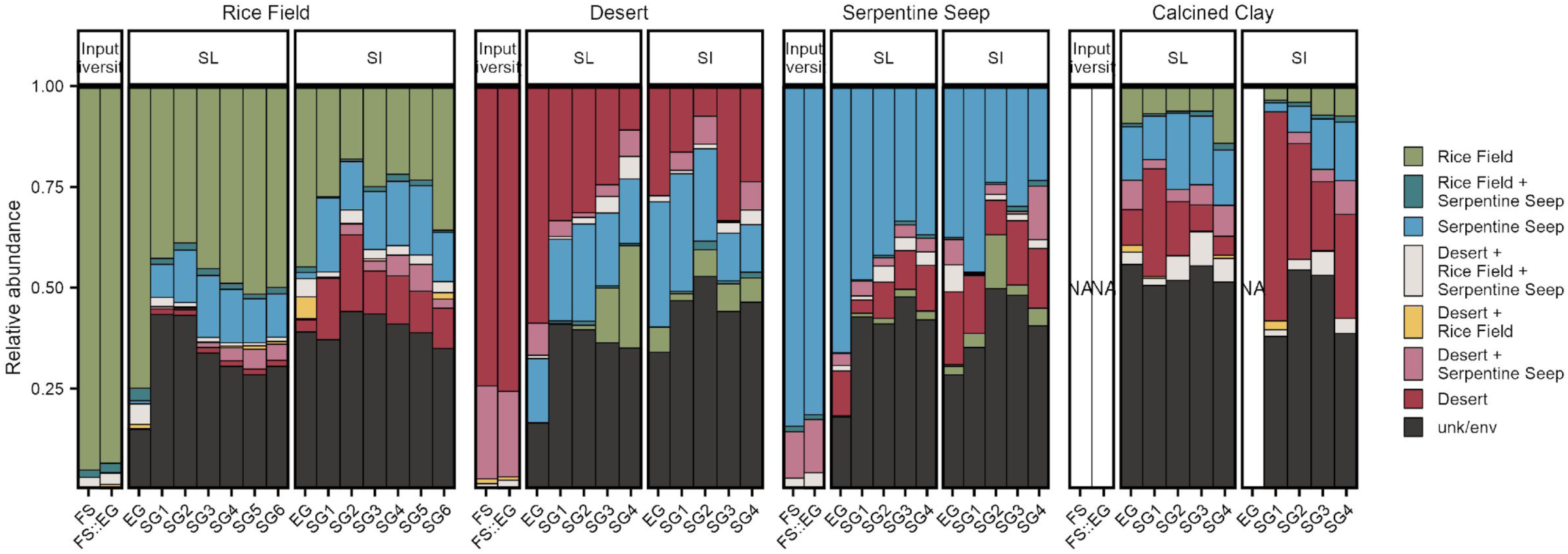
Dispersal homogenizes microbiomes over time. Using both original field soils (FS) and enrichment generation inocula (FS::EG, field soil mixed with sterile calcined clay), we assigned ASVs as originally belonging to one or multiple soil origins and is indicated by color. For example, if an ASV was observed in field soil or inocula samples for Rice Field, but not Desert or Serpentine Seep, it was designated a Rice Field ASV; if observed in Rice Field and Serpentine Seep, but not Desert, it was designated a “Rice Field + Serpentine Seep” ASV. ASVs that were not observed in either the field soil samples or source inocula were assigned “unk/env” to indicate their origin as unknown, but likely from the growth chamber environment. Notably, sterile-inoculated (SI) lines of each soil treatment—which acquired microbiomes from the environment—were each biased towards their respective soil treatments. Calcined Clay, which received no initial inoculum in the enrichment generation was more evenly represented by ASVs from other soil treatments.

Because dispersing taxa enjoyed a fitness advantage within our selection scheme, we wondered if other general bacterial traits, like growth rate, similarly influenced selection. To test whether microbiomes became biased towards fast-growing taxa over time, we estimated the mean doubling time of ASVs observed in SL and SI plants at each generation. Specifically, we aligned our 16S rRNA amplicon sequences to the Estimated Growth rates from gRodon Online (EGGO) database, which includes predicted growth rates for more than 200,000 bacteria and archaea, and assigned a doubling time to each ASV in our Rice Field dataset (Weissman, Hou, and Fuhrman 2021). While we found sterile-inoculated microbiomes to be biased towards fast-growing taxa in all generations, selection line microbiomes became biased toward fast-growing taxa over time (**Figure 6**). Though this analysis offers a coarse view of average growth rates within communities, the bias towards fast-growing taxa in SI, compared to SL microbiomes, is consistent with the fact that rapid colonization of unoccupied space—i.e. SI roots at the start of each generation—often results in a fitness advantage (López et al. 2023; Debray et al. 2022).

**Figure 6.**
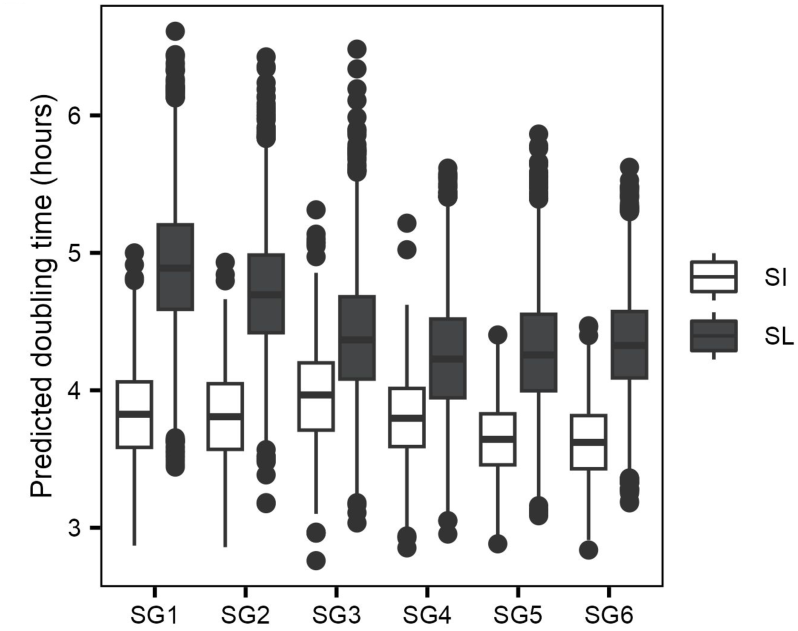
Microbiomes become overrepresented by fast-growing taxa over time. We determined ASV doubling times by matching ASVs to the Estimated Growth rates from gRodon Online (EGGO) database. We then performed 1000 bootstraps randomly subsetting 100 ASVs that were present in each generation for selection line (SL) and sterile-inoculated (SI) microbiomes. Each point represents the mean predicted doubling time of the ASV subset in each bootstrap. By setting generation as a numeric factor and performing OLS regression, we found that predicted doubling time significantly decreased across selection generations in both SL (*t* = -90.9, *p*<2e-16) and SI (*t* = -19.6, *p*<2e-16) samples.

### Selection strength diminishes over time

Having observed the effects of individual selective forces on community composition, e.g. strong environmental and host-mediated filtering (**Figure 4**) as well as abundance patterns influenced by dispersal and growth rate (**Figures 5-6**), we sought to better their overall contribution to microbiome assembly through time. To do so, we determined the extent to which Rice Field selection line microbiomes deviated from a neutral community model in each generation (**Supplemental Figure 7**). A neutral community model assumes that microbiome assembly is the result of stochastic dispersal and drift, and that all individuals have equal fitness; accordingly, deviations from neutrality indicate that community assembly was influenced by non-neutral processes, e.g. active dispersal or differential fitness (Sloan et al. 2006). For each generation, we used abundance data from the previous generation as our source community to parameterize migration and replacement rates and confirmed that this step improved model fit compared to random subsampling as in Burns et al. 2016 (Burns et al. 2016). Assessing the overall fit of the neutral models via generalized R^2^, we found that model fit improved across selection generations (SG1: R^2^=0.109; SG6: R^2^=0.558; **Supplemental Figure 7**). This result suggests that host-mediated selection had a strong influence on microbiome assembly in early selection generations and increasingly stabilized communities. However, because neutral model fit was still poor in SG6, non-neutral processes were still actively determining microbiome assembly in late selection generations.

### Identification of phenotype-associated taxa

Using Rice Field microbiomes, we followed several lines of inquiry that identified a small subset of taxa most likely responsible for improvements in host phenotypes. We used Rice Field data for these analyses because, for logistical reasons, we focused our resources on this soil treatment by performing an additional two selection generations and sequencing all root-associated samples (*n*=1019). Our first two analyses were straightforward, simply asking 1) which ASVs were most abundant in the final selection generation and 2) which ASVs were differentially abundant in droughted versus watered microbiomes. To answer the first question, we ranked ASVs by their mean relative abundance in SG6 and found that one in particular, a species of *Ideonella*, was the dominant taxon across all selection lines (mean relative abundance = 33%) as well as sterile- (25%) and live-inoculated (21%) lines (**Supplemental Figure 8**). Combining this *Ideonella* with the next top ten most abundant ASVs accounted for more than half of SL microbiome relative abundance in SG6 (60%), and nearly half in SI (45%) and LI (47%) microbiomes (**Supplemental Figure 8**). In total, these top 11 ASVs were represented by five Proteobacteria (from the genera *Ideonella*, *Azospirillum*, *Asticcacaulis*, and two within the family *Rhizobiaceae*), two Actinobacteriota (representing *Streptomyces* and *Nocardioides*), and one each from the phyla Bacteroidota (*Flavisolibacter*), Fibrobacterota (*Fibrobacteraceae*), Myxococcota (*Kofleraceae,* likely *Haliangium*), and Verrucomicrobiota (possible *Chthoniobacteraceae* or *Terrimicrobiaceae*).

To answer our second question—which ASVs were enriched in either drought or watered conditions—we measured the differences and effect size of median CLR-transformed abundances of ASVs between selection line (droughted) and live-inoculated (watered) conditions. We then stringently filtered data to only keep ASVs where enrichment in one condition or another was significant after FDR correction, with effect sizes greater than +/- 0.5, and having met these criteria in at least three selection generations. Those remaining included 30 ASVs identified as drought-enriched, as well as 30 ASVs identified as water-enriched (**Supplemental Figure 9**). Of note, Actinobacteriota made up an outsized proportion of drought indicators given the total diversity of Actinobacteriota ASVs observed (*p*<0.05, one-sided Fisher’s exact test, **Supplemental Figure 9**); by contrast Proteobacteria were the most represented phylum among both drought and water indicators, though this group was also the most diverse throughout the experiment (**Supplemental Figure 9, Figure 7A**). Overall, water indicators were more diverse at the phylum level (**Supplemental Figure 9**).

**Figure 7.**
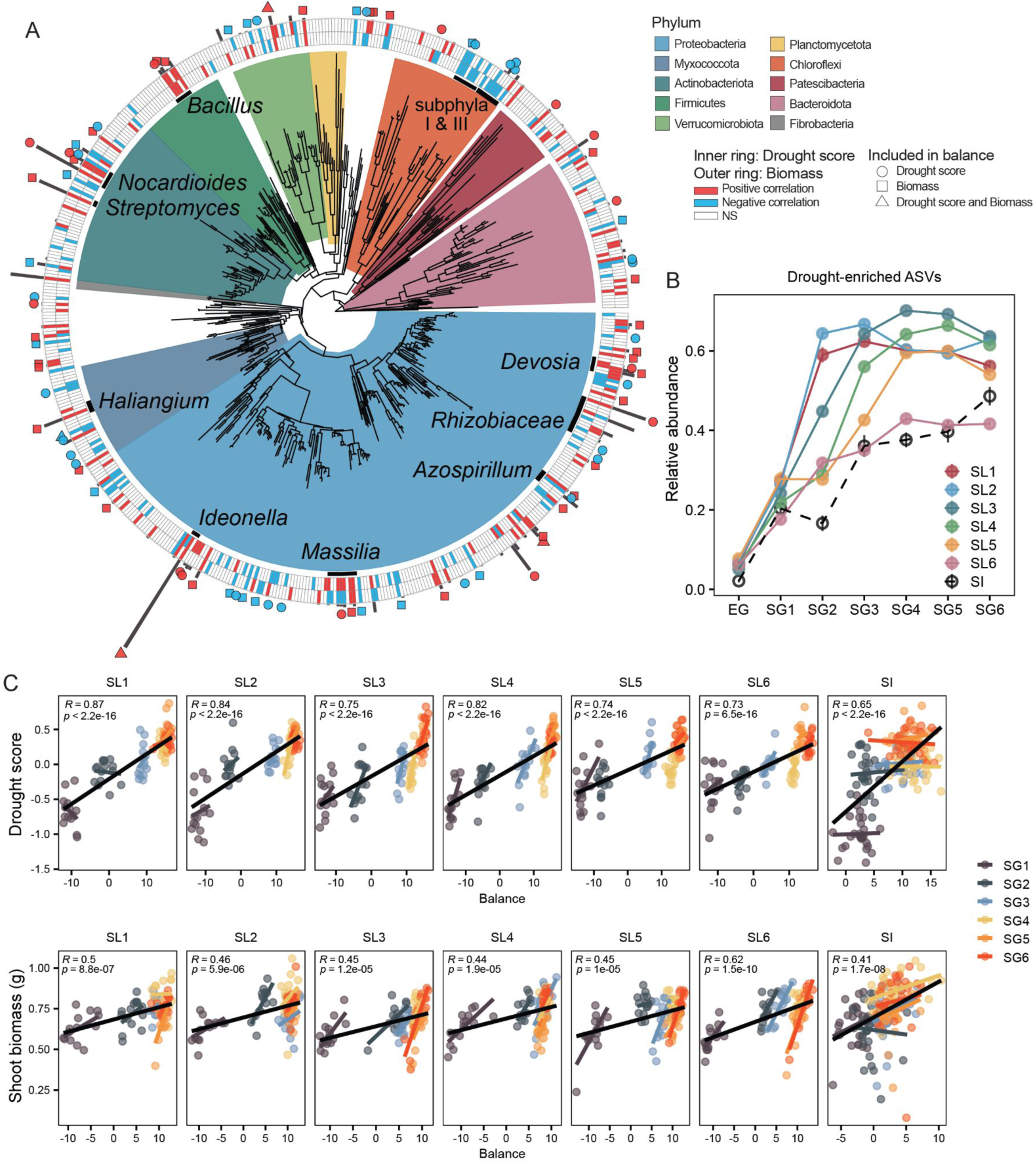
Identifying phenotype-associated ASVs. (**A**) A phylogeny of Rice Field ASVs remaining after four selection generations with branches colored by phylum; black bars and corresponding labels highlight notable clades at lower taxonomic levels. The inner (drought score) and outer (shoot biomass) tiled rings indicate whether an ASV showed a strong positive (red) or negative (blue) response to phenotype, or lack thereof (white). To determine whether a response was strong or not, we ran two mixed effects linear models for each ASV-phenotype combination. One model included selection line as a random factor, while the other included selection generation as a random factor nested within selection line; doing so allowed us to better hone in on ASVs that were both enriched over time and consistently correlated with phenotypes within selection generations. Responses were considered strong if the t-value resulting from one or both models was two or more standard deviations away from the mean. Gray bars extending outward from the tiled rings indicate the median relative abundance of ASVs in the final selection generation (SG6). Symbols beyond the gray bars denote whether ASVs were included in the numerator (red) or denominator (blue) of the balances shown in panel C. (**B**) The mean, combined relative abundance of drought enriched ASVs over time; error bars (present, but small) are +/- one standard error of the mean. Drought enriched ASVs were identified using the effect and t-test functions in the ALDEx2 R package by comparing CLR-transformed abundances between SL (droughted) and LI (not droughted) plants at each selection generation. (**C**) Balances successfully predict drought scores and biomass phenotypes. In brief, balances are compositionally-aware log contrasts in which the numerator and denominator represent two sets of ASVs whose ratio can be used to predict a continuous phenotype <u>(Rivera-Pinto et al. 2018)</u>. We composed distinct balances for each phenotype and performed OLS regressions on individual lines; in each case, SI lines included, balances were significant predictors of plant phenotype. Statistics and black trendlines correspond to the overall regression; colored trendlines show the relationship between balance values and phenotype within generations.

Next, we sought to specifically identify ASVs likely driving growth promotion and improved drought scores (**Figure 7**). By mapping which ASVs showed strong positive or negative correlations with plant biomass and drought score onto a phylogenetic tree, it became clear that though these taxa spanned most of the tree—including many major phyla—there were also several notable clusters representative of genus-level clades with consistent responses associated with host phenotype (**Figure 7A**). Further, many of the most abundant taxa in the final selection generation—outlined above—also strongly correlated with enhanced rice drought tolerance. Notably, multiple *Bacillus* ASVs also exhibited strong association with improved rice drought performance, despite their lower relative abundances in SG6 (ranging from 0.50-0.67%, **Figure 7A**). ASVs identified as belonging to subphyla I and III of the Chloroflexi stood out as strongly negatively correlated with both drought score and biomass phenotypes (**Figure 7A**).

We then used the correlation data to recombine ASVs into balances to find subcompositions of ASVs predictive of phenotypes (**Figure 7C**). In brief, balances acknowledge the compositional structure of microbiome data and that information is therefore carried in the ratios between components (e.g. ASVs); balances, then, are a ratio of the geometric means of two groups of taxa which can be used to predict a continuous phenotype (Rivera-Pinto et al. 2018; Morton et al. 2017). We constructed balances separately for drought score and shoot biomass phenotypes, where the numerator and denominator groups included taxa positively and negatively associated with phenotype, respectively. In total, the drought score balance included 32 ASVs (numerator = 17, denominator = 25) and the shoot biomass balance included 53 ASVs (numerator = 36, denominator = 17). Both balances showed exceptionally strong correlations to phenotypes, with Pearson correlation coefficients as high as 0.87 and 0.62 for droughts score and shoot biomass, respectively (**Figure 7C**). Though balances were constructed from selection line data only, they performed nearly as well applied to sterile-inoculated sample data (**Figure 7C**; drought score: *R*=0.65; shoot biomass: *R*=0.41)

### Functional enrichments among phenotype-associated taxa

Finally, we sought to identify gene functions enriched among our selected taxa in order to better understand the forces potentially influencing their selection during the experiment. To do so, we conducted shotgun metagenomics on a subset of samples from Rice Field selection line plants across all 7 generations (n=37); from this data, we resolved 272 high quality metagenome-assembled genomes (MAGs). To allow for correlations between functional capacity in our metagenomic dataset and the much greater temporal and phenotypic resolution afforded us by our 16S dataset, we assigned individual MAGs to ASVs from the 16S dataset using a combination of taxonomic and abundance data. Next, we identified Clusters Of Orthologous Genes (COG) high-level functional categories with significant enrichments in gene counts among phenotype-associated taxa; specifically, we ran two parallel analyses using the taxa included in the numerators of each of the balances described above (**Figure 8**). As background for each analysis, we used all ASV-matched MAGs (n=165). After applying FDR corrections to p-values and filtering for significance (p<0.05) we found that the groups of MAGs associated with both balances were significantly enriched for genes within the functional categories *Carbohydrate transport and metabolism* and *Transcription* compared to non-selected MAGs. Notably, there were also differences between enrichment profiles for the drought score and shoot biomass MAG sets; specifically, MAGs from the drought score balance showed significant enrichment of genes related to *Cell motility*, *Inorganic ion transport and metabolism*, and *Intracellular trafficking, secretion, and vesicular transport*, while the biomass-related balance MAGs showed significant enrichment of *Lipid transport and metabolism* and *Secondary metabolite biosynthesis, transport and catabolism*.

**Figure 8.**
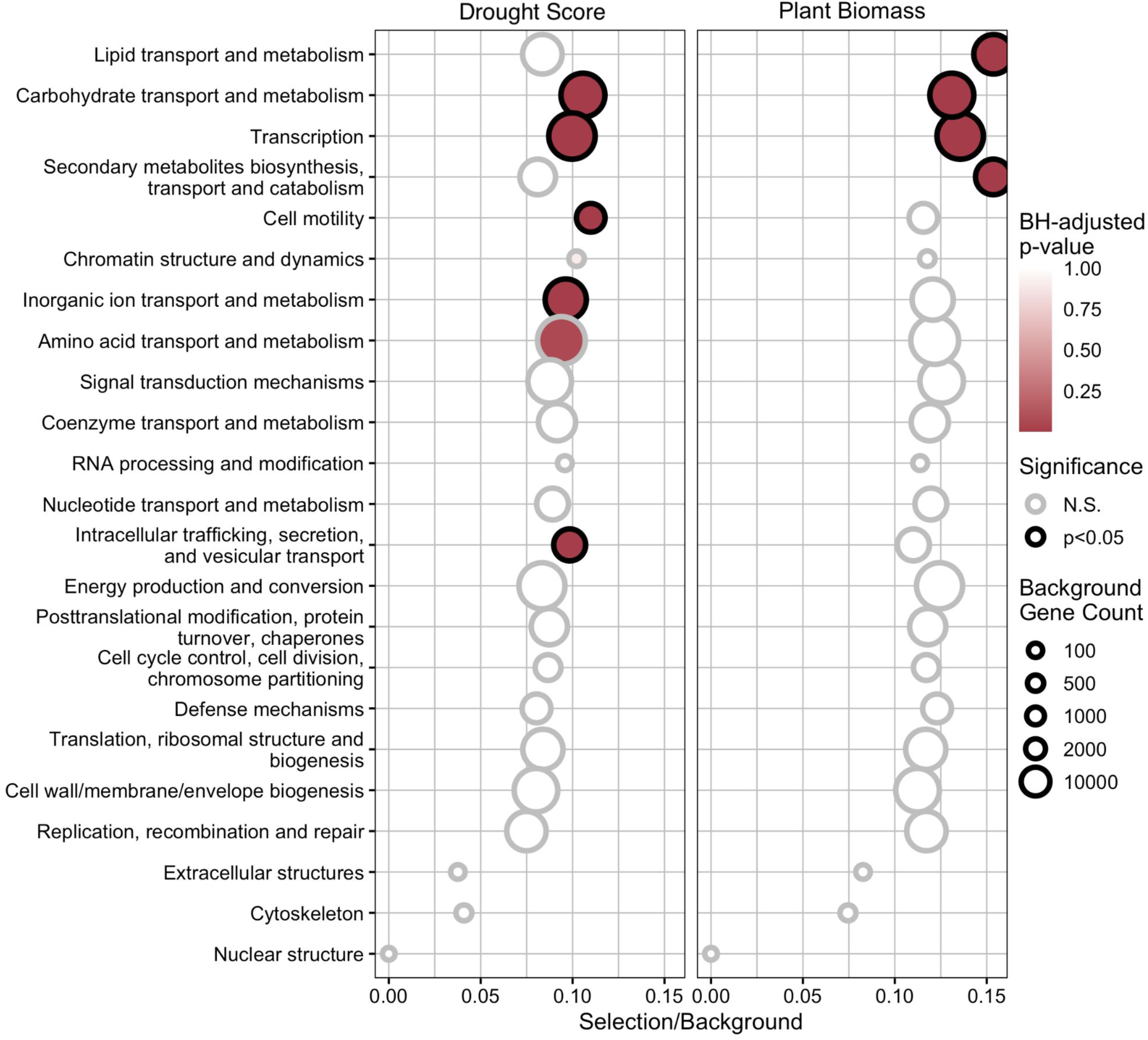
Bacterial communities associated with increased plant biomass and plant vigor during drought are enriched for both common and distinct functions. By comparing the metagenome-assembled genomes (MAGs) of taxa from balance group numerators with all other MAGs in the dataset, we identified high level functional categories that were significantly enriched in the genomes of these groups. Selection/Background refers to the ratio of gene counts among phenotype-associated taxa relative to all taxa for each functional category. The size of points indicates the background gene count, or the total number of genes within a functional category represented by all MAGs. We determined p-values with hypergeometric tests and applied multiple testing correction before determining significance. Functional categories that were significantly enriched in balance numerator groups were then further investigated to identify significantly enriched individual COGs (Supplemental Figures 10-11)

Within these significantly enriched high-level COG functional categories associated with each group, we repeated the same processes above to identify individual COGs with enriched gene counts. Taxa associated with improved drought score showed specific enrichment of genes related to iron transport (COG1629), flagellar structure (COG1344), as well as a diverse array of COGs related to sugar transport and metabolism (**Supplemental Figure 10**).

Interestingly, both groups showed significant enrichment for COGs associated with glycerol-3-phosphate transport (COG1653, COG0395, **Supplemental Figure 11**), a bacterial function identified in recent studies as correlated with drought tolerance in rhizosphere communities (L. Xu et al. 2018). Together, these results suggest that bacteria selected in our experiment are enriched for specific microbial functions associated with transport of a range of metabolic products and nutrients compared to other non-selected taxa in our system.

## Discussion

Here, we present a multi-generation experiment in which we applied host-mediated selection to diverse sources of inocula to derive simplified soil communities that promote growth and drought tolerance in rice. One of our primary motivations for this experiment was to determine the degree to which inoculum source matters, or if any microbial community can serve as a suitable substrate for host-mediated selection. Though microbial communities adapt to their local environments and are therefore biased towards specific taxa and functions (Kraemer and Boynton 2017; Bokulich et al. 2014; Fierer and Jackson 2006; Bååth 1996), we reasoned that any sufficiently diverse community would include members capable of inducing a targeted host phenotype (Delgado-Baquerizo et al. 2016). These members could then be maintained or enriched via host-mediated selection if selection were sufficiently strong relative to stochastic effects or other factors influencing microbial fitness (Belotte et al. 2003; Kraemer and Kassen 2015). Here, however, soil treatments differed in their response to host-mediated selection, where Rice Field and Desert soil microbiota successfully improved rice phenotypes over time but Serpentine Seep did not (**Figure 3**). Interestingly, selection maintained native diversity in Rice Field microbiomes, but preferentially enriched bacteria acquired via immigration in Desert ones (**Figure 5**). We suspect that Rice Field better maintained diversity because its source inoculum likely included more taxa that are better adapted to flooded conditions and capable of colonizing rice roots. Given that plant hosts both actively and passively influence microbial colonization (Zhang et al. 2023; B. Wang and Sugiyama 2020; Johnson et al. 2010), the latter may have been of particular importance because roots—but not soil—were harvested from EG plants to inoculate SG1. However, even Calcined Clay, which relied solely on immigration for microbiome assembly in EG, successfully responded to host-mediated selection. Taken together, these results suggest that source inoculum provenance strongly influenced the trajectory of host-mediated selection, but was just one of many factors determining its outcome.

Dispersal also played an evident role in microbiome assembly in all soil treatments. Despite negligible overlap in shared species at the start of the experiment, all soil treatments generally converged toward a common set of taxa rather than solving the phenotypic “problem” with unique solutions (**Figure 4C, Supplemental Figures 5-6**). Consistent with this observation, others have shown that even low rates of dispersal can homogenize a metacommunity (Fodelianakis et al. 2019). In particular, when factors determining bacterial fitness—i.e. environmental conditions, plant host genotype, and artificial selection criteria—are consistent throughout the metacommunity, dispersal has an even greater homogenizing effect (Evans, Martiny, and Allison 2017; Bell 2010; Kraemer and Kassen 2015). Using the Desert soil treatment as an example, immigrating taxa appeared to enjoy a fitness advantage over the native diversity of the field soil within the context of host-mediated selection; as a result, these taxa were selected and further enriched over time, increasing the compositional similarity between Desert and other soil treatment microbiomes.

As microbiome compositions changed across selection generations, we observed clear patterns of maintenance and enrichment of taxa positively correlated with the rice phenotypes under artificial selection. Many of these taxa are members of genera that include well-known plant growth promoting rhizobacteria (PGPR), including *Azospirillum* (Proteobacteria) and *Bacillus* (Firmicutes), both of which have previously been shown to improve rice growth and drought tolerance (Fukami, Cerezini, and Hungria 2018; Ruíz-Sánchez et al. 2011; Sansinenea 2019; Karmakar et al. 2021). Actinobacteriota, particularly *Streptomyces* and *Nocardioides*, increased in relative abundance over time as well, but were also consistently enriched in droughted (SL) compared to watered (LI) microbiomes (**Supplemental Figure 9**). Many other studies have similarly reported the enrichment of root-associated Actinobacteriota during drought (Naylor et al. 2017; L. Xu and Coleman-Derr 2019; Santos-Medellín et al. 2017; Fitzpatrick et al. 2018) and have also demonstrated *Streptomyces* capable of inducing drought tolerance in grasses, including rice, through osmolytic balancing, reduction of ROS, and auxin production (S. Niu et al. 2022; Selim et al. 2019; Yandigeri et al. 2012).

We also identified taxa that strongly correlated with improved rice drought tolerance, but for which little is known about their potential for plant growth promotion. For example, we observed the enrichment of two Proteobacterial genera, *Devosia* and *Ideonella*, whose abundances strongly correlated with enhanced drought score and biomass phenotypes. While there appeared to be multiple beneficial strains of *Devosia* enriched through host-mediated selection, a single *Ideonella* ASV reached exceptionally high relative abundances in SG6 Rice Field microbiomes (**Supplemental Figure 8**). Isolates described for both genera appear to have several characteristics in common, including production of siderophores and auxin, as well as the ability to fix atmospheric nitrogen (Rivas et al. 2003; Chhetri, Kim, Kang, Kim, et al. 2022; Chhetri, Kim, Kang, So, et al. 2022). Though literature describing interactions of *Devosia* and *Ideonella* with plants is sparse, *Devosia* has been shown capable of nodulating an aquatic legume (Rivas et al. 2003), and members of *Ideonella* can promote the growth of eucalyptus seedlings (De Sousa Fabiano Gama et al. 2022). Overall, our ability to identify PGPR—ranging from well-known to poorly characterized—suggests that host-mediated selection is not only useful for microbiome engineering, but can also be used to identify novel species of plant-beneficial microbes.

Additionally, we identified balances, or subcompositions of taxa, that significantly improved predictions of rice phenotypes compared to the abundance data of individual taxa alone (**Figure 7**). Interestingly, the balances—constructed separately for drought score and shoot biomass phenotypes—showed little overlap with one another, despite the fact that many of the ASVs included in the balances were strongly correlated with both phenotypes (**Figure 7A**). This suggests that discrete subcompositions of the microbiome were important for determining drought score and biomass, separately. While balance numerator groups represent a short list of potential PGPR and candidates for syncom construction, denominator groups represent taxa more likely to hinder plant phenotypes with increased abundance. Though balances were constructed from Rice Field selection line data, they may help to explain the failure of Serpentine Seep to respond to host-mediated selection: Chloroflexi were overrepresented in the denominator group for drought score and was also the phylum designation of the most abundant ASV in final generation Serpentine Seep plants. Balances also corroborate that phenotypic improvements observed across selection generations in sterile-inoculated plants corresponded to similar changes in microbiome composition experienced by their selection line counterparts.

By comparing metagenome-assembled genomes (MAGs) associated with our selected taxa (balance numerators) to all other MAGs in the dataset, we identified specific functions enriched in selected taxa that could help to explain their positive correlation with phenotypes.

Supporting the hypothesis that balance groups had discrete effects on either drought score or shoot biomass phenotypes, few enriched functions were shared between the two groups. This result is perhaps not unexpected given that the two balances were composed of different sets of taxa, but it does enable a closer look at which functions were important for each phenotype individually. For example, genes related to both iron transport (COG1629) and carbonic anhydrase (COG3338) were significantly enriched by the numerator group for drought score but not shoot biomass (**Supplemental Figure 10**). Previous work has specifically demonstrated iron transport and metabolism as important functions for Actinobacterial-mediated drought tolerance in sorghum (L. Xu et al. 2021) and carbonic anhydrase as a potential mechanism for *Bacillus*-mediated drought tolerance in wheat (Aslam et al. 2018). By comparison, genes associated with both lipid and secondary metabolite transport showed strong enrichment within MAGs belonging to the numerator group for biomass. Lipids have been identified as a key avenue of communication between hosts and microbes during plant microbe interactions, including beneficial symbioses (Siebers et al. 2016). Additionally, on the list of significantly enriched secondary metabolite COGs are cytochrome P450s, which are capable of producing a wide array of many small molecules, including phytohormones and other compounds used by microbial partners to manipulate plant fitness (Orozco-Mosqueda, Santoyo, and Glick 2023).

Interestingly, both drought score and biomass balances showed strong enrichment for COGs associated with G3P transport and metabolism (COG1653, COG0395). Prior work in another plant system (*Sorghum bicolor*) has found a correlation between G3P transport in Actinobacteriota and their tolerance to drought stress within the rhizosphere environment (L. Xu et al. 2018); it was hypothesized that this correlation may be due to significant increases in G3P production observed in plants exposed to drought stress (L. Xu and Coleman-Derr 2019).

Interestingly, our data suggest that transport of G3P may also be an important function in drought-related contexts outside the Actinobacteriota, as G3P transport-related genes were present and enriched across multiple phyla represented in our balance numerator taxa, including Actinobacteriota as well as Armatimonadota, Bacteroidota, Firmicutes, Myxococcota, Proteobacteria, and Verrucomicrobiota. Furthermore, the fact that our MAGs were selected based on their associations with hosts exhibiting enhanced drought tolerance, rather than on their own ability to survive drought, suggests that G3P metabolism and transport may play a specific role in meditating mutualistic interactions between plants and microbes under water stress.

Overall, this study demonstrates that host-mediated selection efficiently operates at the community level to reduce microbiome complexity and enrich functions of interest. In the context of the nascent body of host-mediated selection literature, it also provides further insight into numerous aspects of experimental design that can present challenges to successful community selection. For instance, the use of appropriate control lines is prudent, but difficult to implement. Specifically, in our study, dispersal complicated our efforts to utilize the sterile-inoculated (SI) lines as meaningful controls. Rather than statically recruiting the same environmental microbes, SI plants recruited microbes from an increasingly optimized metacommunity each generation.

Consequently, sterile-inoculated plants saw phenotypic improvements similar to their selection line counterparts. The shared trajectory of selection and intended control lines appears to be a consistent phenomenon—to varying degrees—in other host-mediated selection studies (Panke-Buisse et al. 2015; King et al. 2022; Garcia et al. 2022; Kalachova et al. 2022;

Rodríguez et al. 2023). “Add-back” controls—where hosts are inoculated from a stock source of the original inoculum at the start of each generation—were subject to the same phenomenon, though to a lesser extent compared to sterile-inoculated or “random-selection” controls (King et al. 2022; Garcia et al. 2022). The use of fully gnotobiotic plants could eliminate this issue, but would be both challenging and expensive to implement for most plant species.

Additionally the length of experimental generations influences whether slow-growing microbial species are maintained in selected communities and determines the likelihood of community turnover within a generation (Wright, Gibson, and Christie-Oleza 2019). Short generation times correspond to high dilution rates (i.e. how frequently microbiomes are selected) and can result in dilution to extinction (Adamberg and Adamberg 2018; Díaz-García et al. 2021; Xie and Shou 2021). Despite our efforts to make generations as long as possible (harvesting plants after 40 days of vegetative growth, before source-sink relationships begin to change with the onset of heading), we still observed selection biased towards fast-growing taxa (**Figure 6**). This may be due in part to the fact that rapid growth can also provide fitness advantages, particularly by influencing priority effects in early stages of microbiome assembly (López et al. 2023; Whitman et al. 2018; Debray et al. 2022). While longer generations may help to maintain slow-growing taxa, they also increase the likelihood of community turnover, which is problematic if taxa driving the phenotype of interest are displaced before they can be transferred to the next generation (Wright, Gibson, and Christie-Oleza 2019). In short, we encourage others attempting host-mediated selection to consider the ways in which experimental design might weaken the strength of artificial selection or otherwise limit the number of potential routes microbiome selection can take.

## Conclusion

Microbiomes have the potential to increase agricultural sustainability and productivity, yet engineering beneficial plant microbiomes remains largely intractable. One of the primary challenges is accurately predicting which microbes will cooperatively improve host phenotype under heterogeneous field conditions. Screening large isolate collections for specific plant growth promoting functions can be a costly and time-consuming process and ultimately fails to account for interactions between microbes, the plant host, and the environment. An approach like host-mediated selection, however, significantly simplifies the process of microbiome engineering by allowing eco-evolutionary processes to optimize microbiome composition at the community level. In other words, host-mediated selection inherently accounts for microbe-microbe interactions (including higher order interactions) while identifying individuals and consortia that are beneficial to plant phenotype. Moreover, selected communities not only provide a blueprint for syncom design but are also a source from which phenotype-promoting taxa can be more readily isolated. From our work, we show that although experimental design must be carefully considered to ensure the greatest chances of success, host-mediated selection holds great promise for identifying and mining plant beneficial taxa from complex soil environments.

## Methods

### Rice growth conditions

We used a near isogenic rice japonica cultivar known as “Super Dwarf” (*Oryza sativa* L.), which is a gibberellic acid mutant that grows vigorously, but only reaches heights of around 20cm at maturity (Frantz and Bugbee 2002). We initially bulked seeds through selfing until we acquired enough material for the entire experiment and stored these at 4°C. Prior to each generation, seeds were surface sterilized then germinated in sealed 50 ml falcon tubes filled with autoclaved milliQ water at 28°C for five days before being transplanted into inoculated substrate. To minimize cross contamination between replicates, each plant was grown in an individual, autoclave-sterilized “conetainer” (Greenhouse Megastore, catalog No. SC10R) and given a separate water reservoir (Restaurant Equipment Solutions, catalog No. LIBB-551HT) (**Supplemental Figure 4A**). We built custom racks to hold five plants each to allow rotation of plants within a soil treatment to normalize environmental and ballast exposure over the course of the generation (**Supplemental Figure 4B**).

The entirety of this experiment was performed in a walk-in growth chamber (**Supplemental Figure 4B**); conditions were set to 16-hour days, during which daytime temperature and humidity were held at 28°C and 80%, respectively, and relaxed to 24°C and 75% humidity at night. Once transplanted, rice were grown for a total of 40 days. We fertilized plants with 10mL 1X Murashige and Skoog medium at the time of planting, and again on days 7 and 14; on day 21 we applied a final fertilization of 10mL 0.5X 20-20-20 NPK. We previously observed that this level of fertilization was growth-limiting but not stress-inducing. Plants were consistently well-watered with milliQ water throughout the enrichment generation; in subsequent selection generations, a drought treatment was initiated after 30 days of well-watered growth.

On day 31, we emptied water reservoirs and allowed the formerly flooded substrate to drain before refilling the reservoir to 25% full capacity; we repeated this emptying and refilling again the following day. On day 33 reservoirs were emptied without refilling and water was withheld for the remainder of the generation.

### Drought score measurements

To non-destructively phenotype plants in high throughput, we built custom hardware and computational scripts to measure Normalized Difference Vegetation Index (NDVI) of individual plants longitudinally. Originally developed for remote sensing applications, NDVI is commonly used to track plant stress status (Rouse et al. 1974; Huang et al. 2021). It exploits the tendency of plants to reflect wavelengths of far and near-infrared light regardless of stress status while increasingly reflecting wavelengths of visible light as stress becomes more severe (Ustin and Jacquemoud 2020; Schepers et al. 1996). Prior to host-mediated selection, we performed several preliminary experiments to confirm that NDVI values produced by our setup were sensitive to changes in rice drought stress status over time (**Supplemental Figure 2**). We confirmed that NDVI decreased with increasing drought stress and also showed that NDVI highly correlated with rice percent water content (**Supplemental Figure 2D-E**).

To obtain NDVI values for plants, we first imaged them within a plywood lightbox (**Supplemental Figures 2A, 4B**), the interior of which was painted with an ultra light-absorbing black paint Culture Hustle USA, Black 3.0); all exterior corners sealed to ensure no light contamination while full spectrum and far red (730nm) LED strips served as light sources within the box (LEDLightsWorld, full spectrum: <u>HK-CRI95F5050X30-X-6</u>, 730nm: <u>5M-HK-8MM-F2835-735-30-NW-IR-12</u>). Two cameras simultaneously imaged plants, one a standard DSLR (Canon, EOS Rebel T7), the other a DSLR modified to be able to detect red and infrared light (Canon, EOS Rebel T1i) (**Supplemental Figure 2A-B**). Both cameras were set to: shutter speed = 1/40; F-stop = 4.5; ISO = 100; Effect = Neutral; White balance = 7000K; Auto lighting optimizer = OFF. The modified camera had previously been sent to LifePixel (Life Pixel Infrared; Mukilteo, USA) where the factory infrared sensor filter was replaced by a dual-pass filter with high transmission from 400-600nm and 700-800nm and complete out of band blocking. Consequently, both cameras assign green and blue values near-equivalently, but have non-overlapping perceptions of red (**Supplemental Figure 2C**).

Paired images were then passed through a custom pipeline to derive NDVI. In particular, we relied heavily on tools from PlantCV (Fahlgren et al. 2015). All scripts are available on GitHub (github.com/colemanderr-lab/Styer-2024). In brief, we used a small section of black background to standardize all images by subtracting the median RGB values within that window from all pixels. Then, we used naive Bayes classifiers—previously trained on images from each camera—to segment plants from background; individual plants were then identified via object detection in the resultant binary mask and saved as separate images. We then extracted color data from each plant, including median red, green, and blue pixel intensities. Using these data from each set of paired images we calculated NDVI as the sum of median red values for both modified (MOD) and standard (STD) cameras divided by their difference:

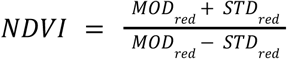

To derive drought scores, we then calculated the area under the curve of NDVI (AUC NDVI) for each plant by fitting a smoothed spline to NDVI values across the 10 days of drought (**Supplemental Figure 2D**); this allowed us to evaluate cumulative drought performance in a single metric. Lastly, because plant biomass was consistently anti-correlated with drought score (**Supplemental Figure 12**), we controlled for its effect by calculating the residuals of the linear model AUC NDVI ∼ shoot biomass. The final drought score, then, represents plant performance across a drought time series independent of the influence of plant biomass.

### Selection scheme

Selections were made independently for each selection line and were determined by both fresh weight and NDVI. As a first filter, microbiomes were only eligible for selection if the fresh weight of their host plant was equal to or greater than the mean of the selection line. We set this initial filter to avoid selecting plants stunted by dysbiosis, having anticipated the negative correlation between NDVI and biomass but not yet having derived biomass-adjusted drought scores; without this initial filtering, we thought it possible, if not likely, selections based on NDVI alone would lead to the accumulation of pathogens. From the microbiomes that passed the biomass-based filter, we selected three from plants with the greatest cumulative NDVI score (i.e. the sum of daily NDVI values); these were then used to inoculate 15 plants to continue the selection line into a new generation. The two next best-performing microbiomes from each selection line were used to create inocula for the subsequent generation SI and LI control lines.

### Inoculum preparation

For the enrichment generation (EG), we prepared inocula simply by mixing equal parts field soil (i.e. Rice Field, Desert, or Serpentine Seep) with sterilized calcined clay by weight. For sterile-inoculated lines, field soils were autoclaved in small batches for 90 minutes prior to mixing to ensure thorough sterilization. Calcined Clay soil treatment plants only received sterilized calcined clay as “inocula”. Pre-germinated rice plants were directly transplanted into the mixed substrates. The goal of the enrichment generation—during which no drought stress was applied—was to first enrich microbes that form associations with rice in non-stressed conditions. Endeavoring to minimize soil edaphic effects and to bias communities towards root-associated bacteria, we only harvested roots and adhering soil—but not bulk soil—from selected EG plants to inoculate the first selection generation (SG1).

To prepare selection generation inocula, we first shook roots to remove loosely adhering soil, finely minced them with a sterile razor, then further macerated the collection with a mortar and pestle. For SG1, the resultant root paste was thoroughly mixed into sterilized calcined clay and rice plants directly transplanted into the inoculated substrate. For SG2 and later selection generations, both root paste *and* bulk soil from selected plants were mixed into sterilized calcined clay prior to rice transplantation. For each soil treatment, microbiomes selected to inoculate SI and LI plants underwent the same preparation with the exception that the fraction earmarked for SI was autoclave-sterilized prior to mixing with calcined clay.

### 16S rRNA library preparation

At harvest, roots were shaken loose of soil, which, for droughted samples, amounted to removing virtually all calcined clay particles. We did not further separate root rhizospheres from endospheres and so we considered microbial community profiles collected from such samples as “root associated.” All roots were manipulated with clean gloves on sheets of single-use aluminum foil; a portion of roots from each harvested plant were immediately placed in 15 ml falcon tubes, flash frozen in liquid nitrogen, and stored at -80°C. We later ground roots with mortar and pestle in liquid nitrogen to prepare samples for DNA extraction. Approximately 0.1 g of the resulting powder was added to Qiagen DNeasy 96 PowerSoil Pro kits and followed all manufacturer’s instructions to extract DNA (Qiagen, Hilden, Germany; catNo: 47017).

Next, we quantified extracted DNA (Invitrogen, Waltham, MA, USA; Quant-iT dsDNA Assay Kit, CatNo. Q33120) and normalized all samples to equal concentrations prior to PCR. All PCRs were performed in triplicate, each reaction consisting of 2uL DNA template at 5 ng/uL, 10uL PlatinumII Hot-Start PCR Master Mix (2x) plus 2uL GC-enhancer (Invitrogen, Waltham, MA, USA; CatNo. 14000014), 0.19uL of 100uM mitochondrial PNA (GGCAAGTGTTCTTCGGA; PNA Bio, Thousand Oaks, CA, USA; CatNo. MP01-50), 0.19uL of 100uM chloroplast PNA (GGCTCAACCCTGGACAG; PNA Bio, Thousand Oaks, CA, USA; CatNo. PP01-50), 0.5uL of 10uM of each forward and reverse primer targeting the V3-V4 region of the 16S rRNA gene, and water to bring final volume to 25uL. From 5’-3’ complete primer sequences included an Illumina adapter sequence, a 12 base pair index, an Illumina TruSeq primer sequence, a 1-7 base pair spacer to increase flow cell diversity, followed by forward (CCTACGGGNBGCASCAG) and reverse (GACTACNVGGGTATCTAATCC) 16S rRNA primer sequences; a complete list of primer sequences is available in **Supplemental Table 2**. Thermocycling conditions were set to follow: an initial 3 min cycle at 94°C, then 36 cycles of 1 min at 94°C, 30 sec at 48°C, 1 min at 72°C, and a final cycle of 10 min at 72°C before holding at 4°C indefinitely. Replicate PCRs were then pooled and quantified before pooling all libraries in equimolar amounts. Pooled libraries were then cleaned with AMPure XP beads (Beckman Coulter, Brea, CA, USA; CatNo. A63880) following the manufacturer’s protocol, eluted in water, then submitted to University of California, Berkeley’s QB3 Genomics for Illumina MiSeq v3 300bp paired-end sequencing (Berkeley, CA, RRID:SCR_022170; Illumina, San Diego, CA, USA).

### 16S rRNA sequence data processing

Demultiplexed reads received from QB3 Genomics were imported into QIIME2 (Bolyen et al. 2019) where primer sequences were removed using the Cutadapt plugin (Martin 2011). Reads were then trimmed to remove low-quality base pair calls before being passed to the DADA2 plugin (Callahan et al. 2016) for denoising and dereplication, resolving sequences into amplicon sequence variants (ASVs). Representative sequences of ASVs were then assigned taxonomies via the feature-classifier plugin (Bokulich et al. 2018) using the sklearn method (Pedregosa et al. 2012) and a classifier trained on the SILVA 138 SSU Ref NR 99 16s rRNA reference database (Quast et al. 2013). Lastly, prior to downstream analyses, we filtered ASVs identified as either mitochondria or chloroplast and additionally removed any ASVs that were observed in only one sample (out of 1425) and with fewer than five reads.

### Shotgun metagenomic sequencing and analysis

Thirty-seven samples were selected for shotgun metagenomic sequencing, each representing a distinct Rice Field selection line at different time points spanning the seven generations of the experiment. DNA was extracted from bulk soil combined from multiple plants and extracted using Qiagen DNeasy 96 PowerSoil Pro kits following all manufacturer’s instructions (Qiagen, Hilden, Germany; catNo: 47017). All samples were then prepared by Novogene (Novogene Corporation Inc., Sacramento, CA, USA) as 150bp paired-end libraries and sequenced the Illumina NovaSeq platform, yielding an average 111 million reads per library. Reads from each library were individually assembled *de novo* using MEGAHIT (D. Li et al. 2015, 2016). Next, reads from each library were aligned to the accompanying assembly via BWA-MEM (H. Li 2013). Three separate binning approaches: CONCOCT (Alneberg et al. 2014), MaxBin 2.0 (Wu, Simmons, and Singer 2016), and MetaBAT2 (Kang et al. 2019) using both covariant tetranucleotide frequencies were applied to each library. Resultant bin sets were combined and refined from these three approaches to produce a single set of high quality bins using DAS Tool (Sieber et al. 2018). Bins from all libraries were subsequently combined and de-replicated with dRep (Olm et al. 2017) to create a final set of metagenome-assembled genomes (MAGs). Removing MAGs with >10% estimated contamination from other taxa and <70% genome completion based on collocated sets of genes that are ubiquitous and single-copy within a phylogenetic lineage (Lee 2019) yielded a final set of 272 high quality MAGs. Each MAG was assigned taxonomic classification using GTDB-tk (Chaumeil et al. 2022). Finally protein coding sequences were identified with prokka (Seemann 2014) and annotated functionally using eggnog-mapper (Cantalapiedra et al. 2021). MAGs were manually assigned to ASVs from the 16S amplicon sequencing analysis using both closest taxonomic matches and comparing ASV and MAG abundances. In total, 165 MAGs were matched to ASVs.

In order to test for microbial functions associated with selected plant phenotypes, we assessed the gene content of ASV-matched MAGs assigned to balance numerators for both drought score and biomass plant phenotypes. We applied the hypergeometric test to identify functional enrichments within these balance numerators relative to the background of all 165 ASV-matched MAGs. In the first round, we tested COG categories (Galperin et al. 2021). Briefly, all genes in the numerator MAGs were grouped and tested for enrichment of genes in each COG category relative to the background of the genes among all 165 ASV-matched MAGs using the hypergeometric test as implemented in ‘fgsea’ (Korotkevich et al. 2016). In the second round, we tested individual COGs among significantly enriched categories identified in the first round.

### Data and code availability

Amplicon and shotgun metagenomic sequence data reported in this study are available in NCBI Bioproject PRJNA1067324. Scripts used to process images and analyze data are available on GitHub (github.com/colemanderr-lab/Styer-2024).

## Acknowledgements

This research was supported by the National Aeronautics and Space Administration under the grant number NNX17AJ31G as well as by the United States Department of Agriculture ARS CRIS Grant 2030-12210-002-00D. This work was performed in part at the University of California Natural Reserve System McLaughlin Natural Reserve DOI: <u>10.21973/N3W08D</u>. We appreciate insights from Ulrich Mueller that helped guide our initial experimental designs and thank Don Brubaker, Michael Derrico, Cathy Koehler, Kent McKenzie, Nicole Tautges, and Carrie Woods for their help facilitating soil collections. We also thank Ciana Caddell, Dawn Chiniquy, Citlali Fonseca-Garcia, Rachel Klaras, Bradie Lee, and Liana Ogden for their help harvesting and processing root samples. Lastly, thank you to Dan Naylor, Jen White, and Edi Wipf who provided helpful comments and suggestions prior to submission.

---

**Supplemental Figure 1.**
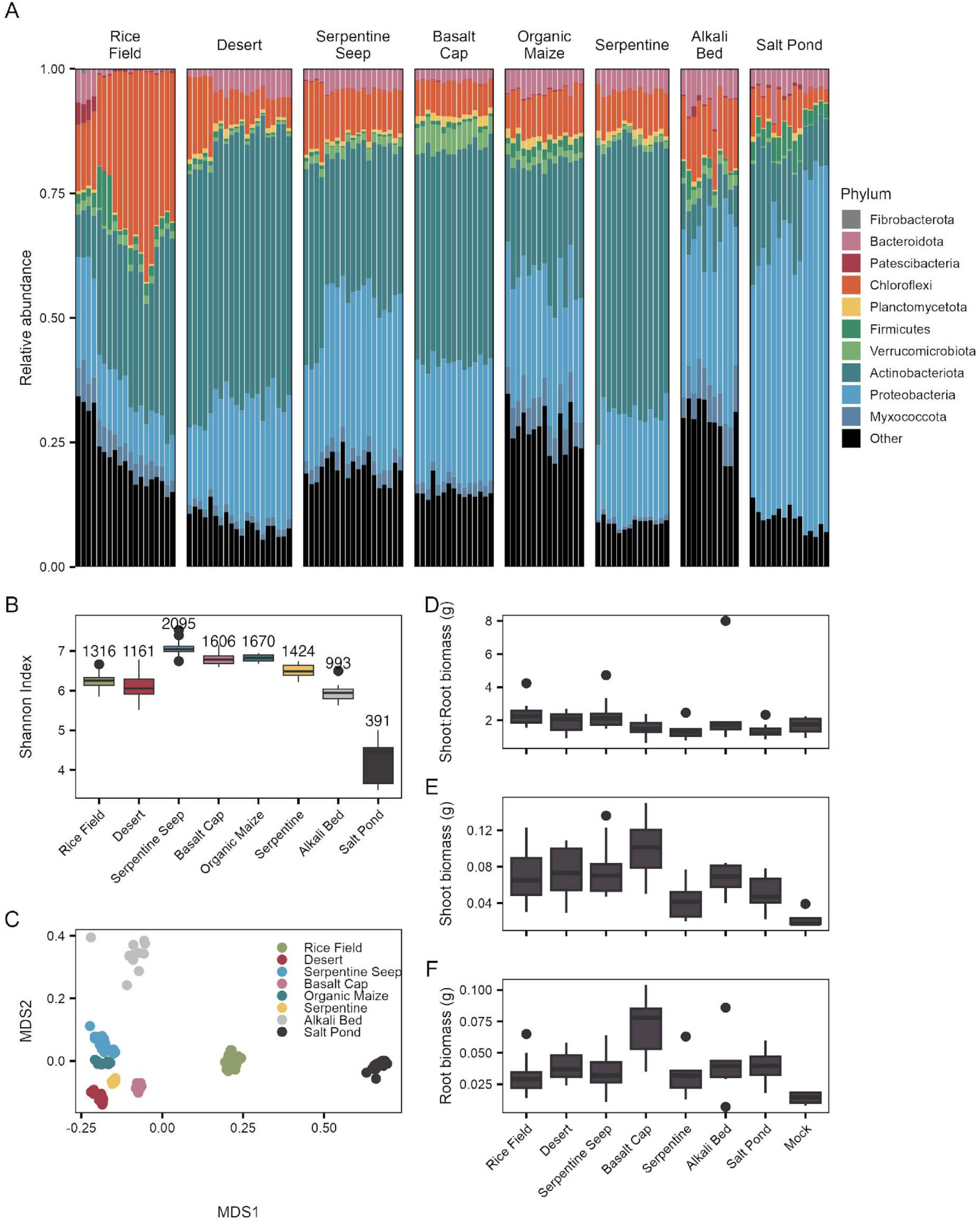
Bacterial diversity of field soils collected as candidate source inocula for host-mediated selection. (**A**) Relative abundance barplots colored by Phylum show large differences between soil bacterial communities even at coarse taxonomic levels. Each bar represents an individual technical replicate. (**B**) Shannon-Index diversity for each soil. The mean number of observed ASVs are shown above each boxplot. (**C**) NMDS-ordinated Bray-Curtis dissimilarity between soils. (D-F) Phenotypic data from a pilot study where rice plants were either inoculated with the field soils shown in A-C or mock inoculated.

**Supplemental Figure 2.**
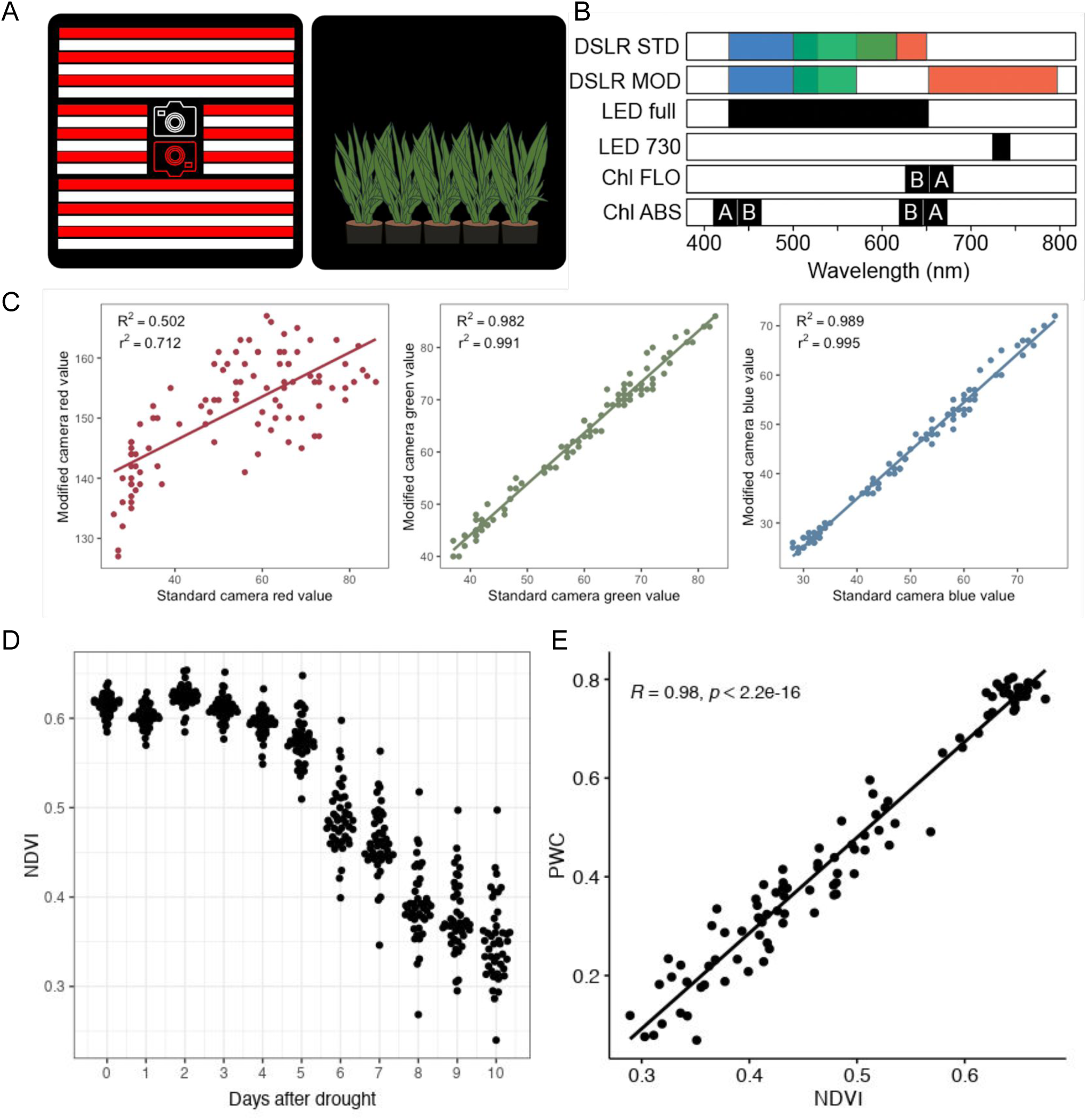
(**A**) Simplified schematic of lightbox design depicting full spectrum (white) and 730nm (red) LED light strips as well as the position of the standard (white) and modified (red) cameras. The interior of the box was painted with an ultra light-absorbing black paint. A shared shutter trigger ensured images from both cameras were captured simultaneously; up to five plants were imaged at the same time. (**B**) A representative drawing of the portions of visible and near-infrared light each camera’s red, green, and blue channels were sensitive to. Beneath, black bars indicate the range of emission spectra of light sources as well as wavelengths expected to be absorbed (ABS) and fluoresced (FLO) by chlorophyll. (**C**) Comparisons of the standard and modified cameras’ median values for red, green, and blue channels individually; each dot represents an individual plant. (**D**) NDVI responds to rice drought stress status over time. Generally, the interval between two time points with the largest difference corresponds to the visible onset of leaf curling symptoms (data not shown). (**E**) At harvest, NDVI is highly predictive of rice percent water content.

**Supplemental Figure 3.**
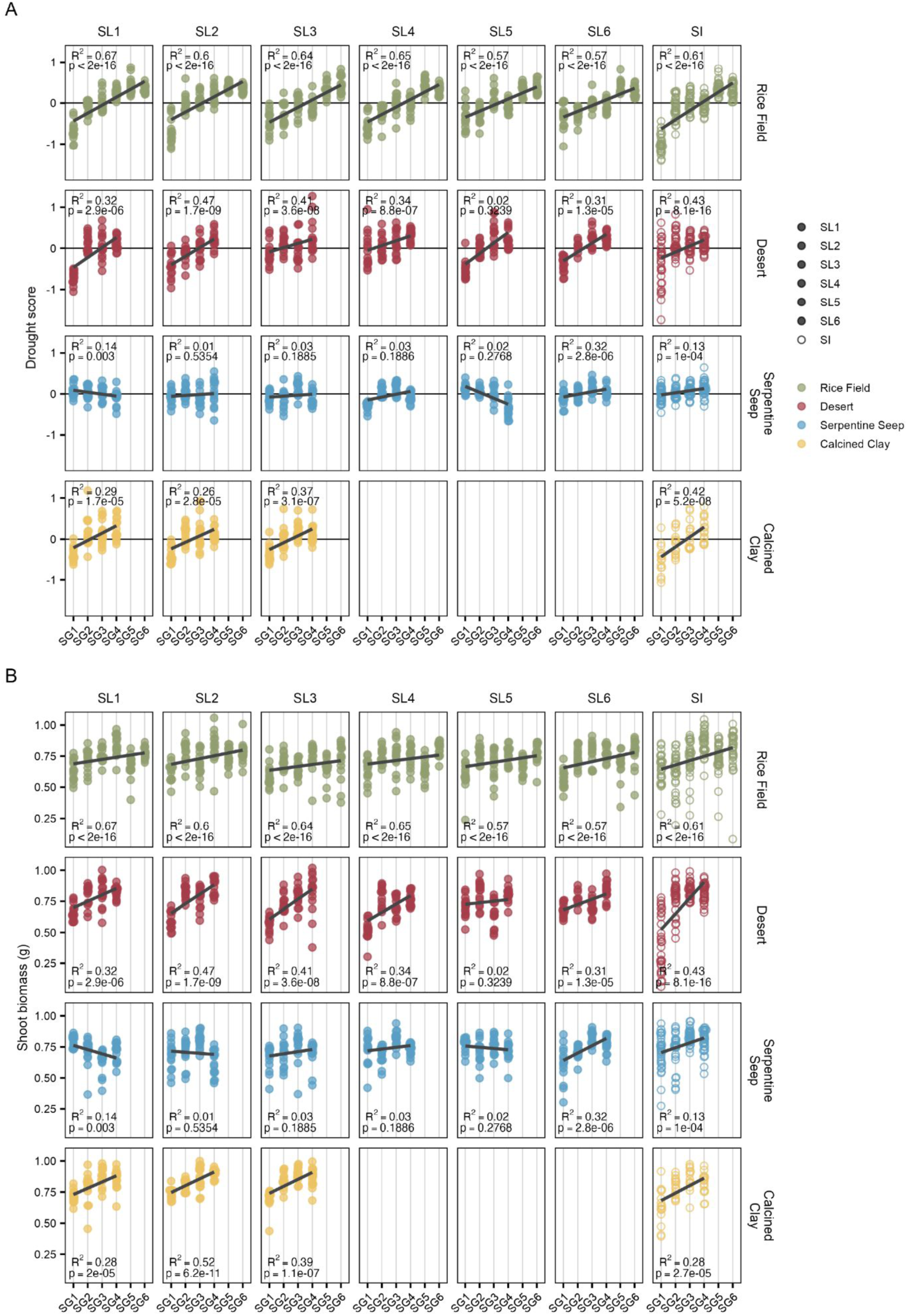
Drought score and biomass phenotypes for individual selection lines. (**A**) Drought scores over time. Statistics and trendlines result from OLS regression. Each point represents an individual plant. Filled circles correspond to selection line (SL) plants; open circles correspond to sterile-inoculated (SI) plants. (**B**) Shoot dry weight biomass over time.

**Supplemental Figure 4.**
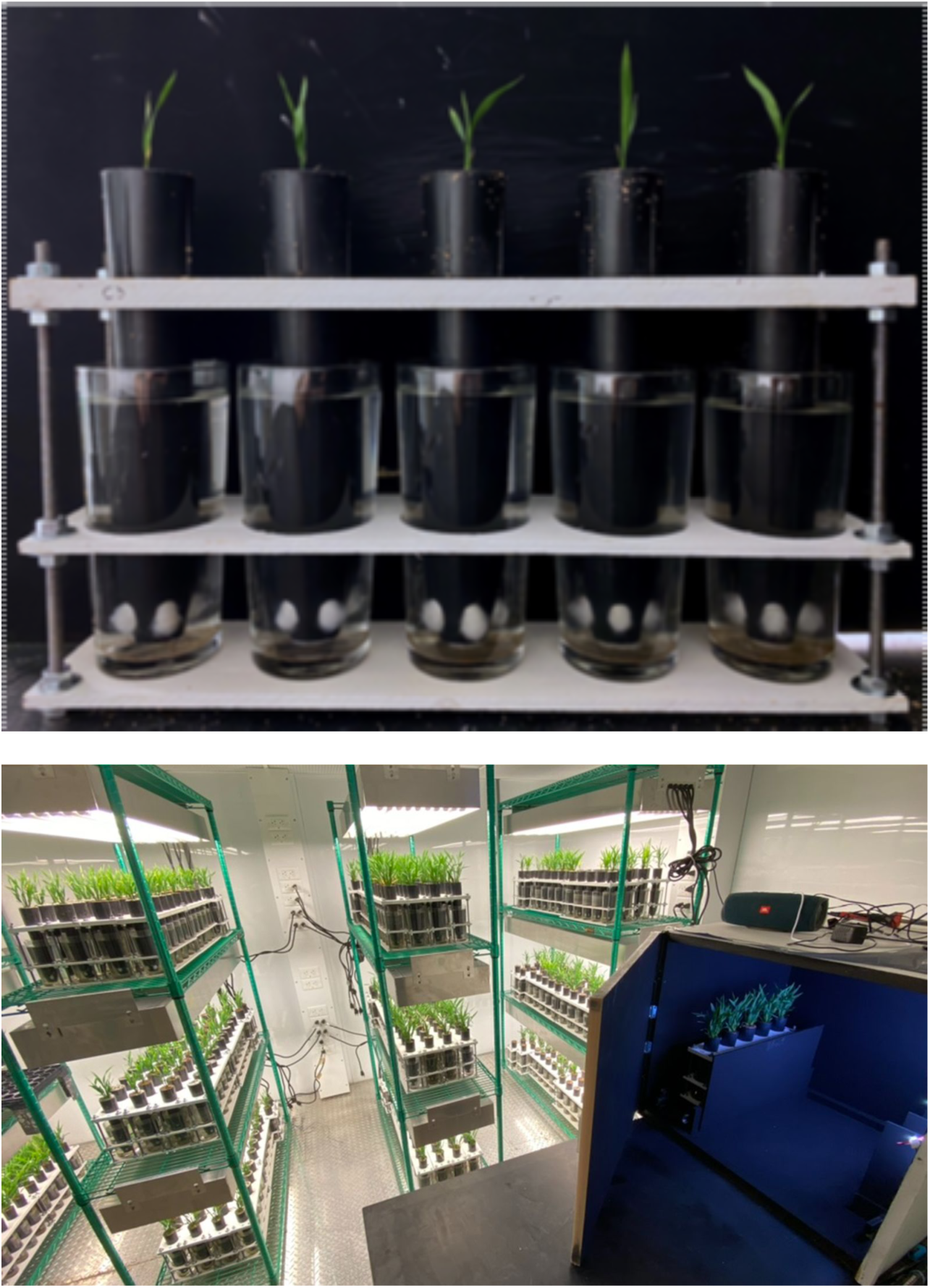
Rice growth conditions. (**A**) Rice were grown in spatially discrete conetainers and water reservoirs to minimize admixture of microbiomes. A small ball of polyester fiber was placed at the bottom of conetainers to prevent substrate from spilling out. Reservoirs were filled with milliQ water which calcined clay substrate was able to wick to the top surface of the container. (**B**) Host-mediated selection was performed in a walk-in growth chamber set to 16-hour days, during which daytime temperature and humidity were held at 28°C and 80%, respectively, and relaxed to 24°C and 75% humidity at night. Each soil treatment was grown on its own shelving unit, but plants were rotated within that unit every three days to ensure each plant experienced the same overall environment and ballast exposure. The lightbox with a rack of plants ready to be imaged can be seen in the bottom right corner.

**Supplemental Figure 5.**
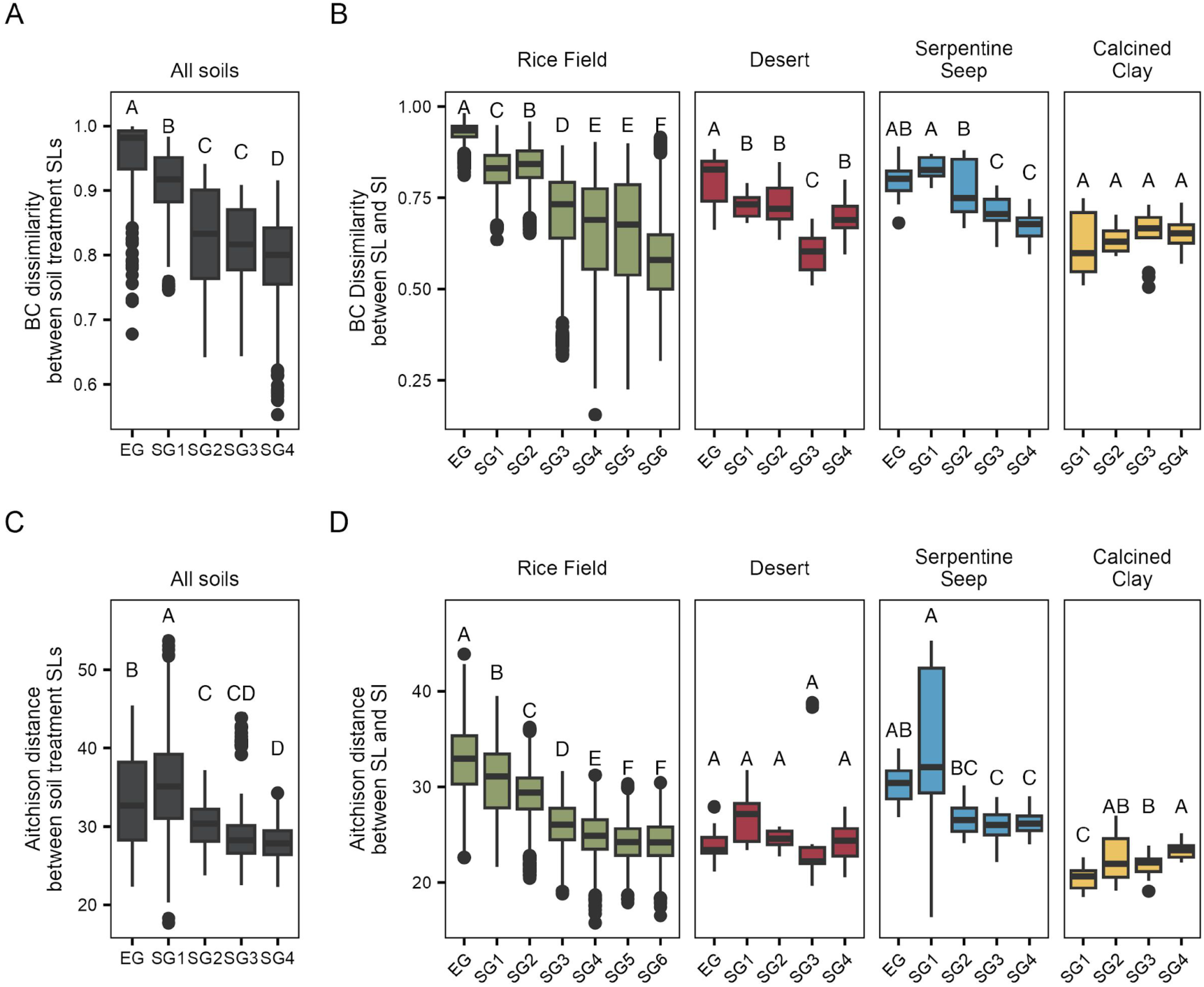
Bray-Curtis dissimilarity (**A-B**) and Aitchison distance (**C-D**) between and within soil treatments across selection generations. Panels A and C show dissimilarity/distances between soil treatment selection lines by BC dissimilarity and Aitchison distance, respectively. Panels B and D are similar, except dissimilarity/distance is between selection line and sterile inoculated microbiomes within soil treatments. BC values are the average of 1000 permutations where each sample was individually subset to 5000 reads. Aitchison distance was calculated as the Euclidean distance between samples after read counts were transformed via robust centered log ratio transformation. Letters above boxplots are the result of a post-hoc Tukey HSD with 95% confidence intervals (*p*<0.05) to detect significant differences between groups. Rice Field samples were subset to match the sequencing effort of other soil treatments when generating data for panels A and C.

**Supplemental Figure 6.**
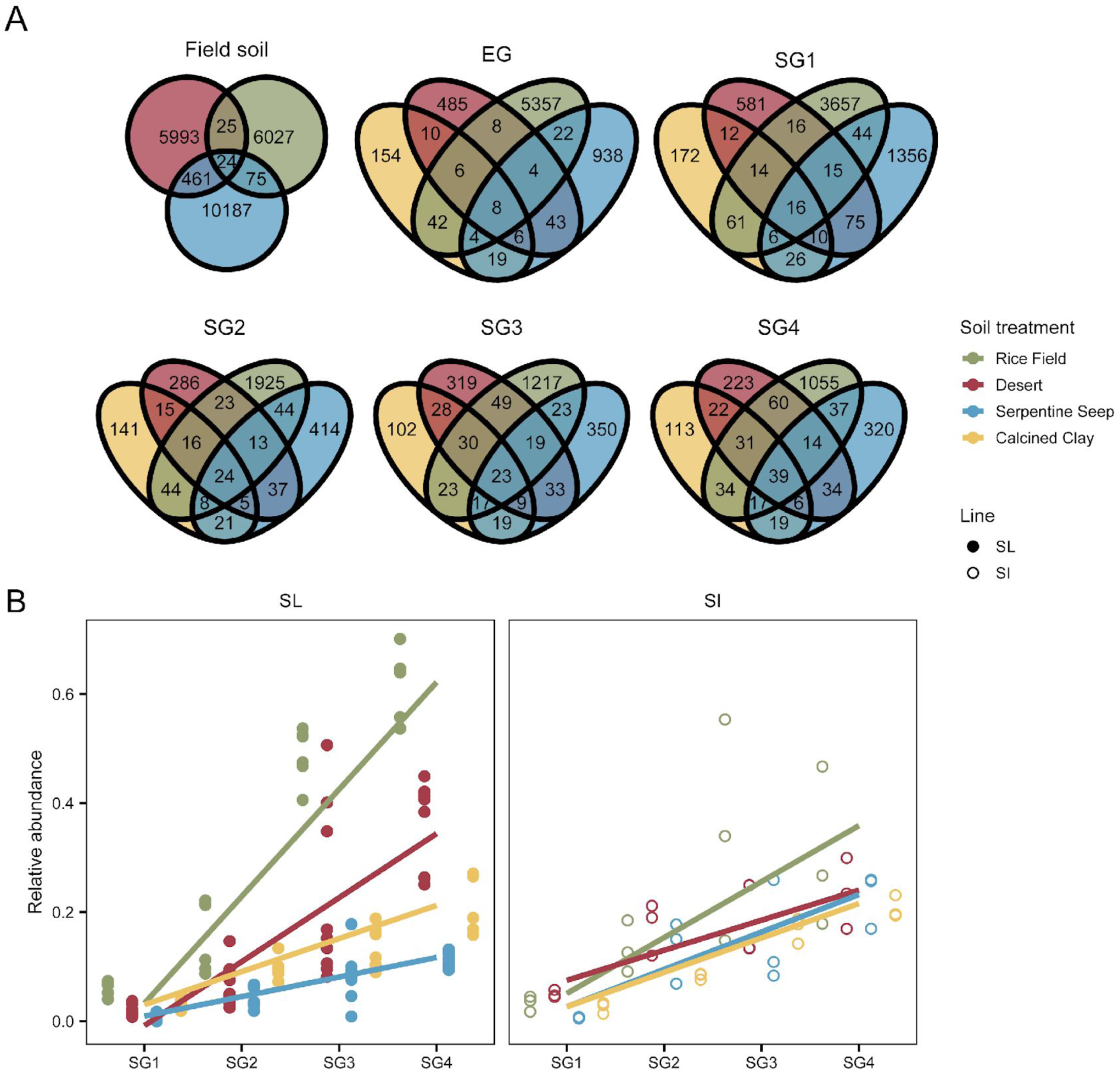
The number and abundance of ASVs shared between soil treatments increased over time. (**A**) Venn diagrams for the numbers of ASVs shared between soil treatments at each generation. Venn components are colored by soil treatment: Rice Field (green), Desert (red), Serpentine Seep (blue), and Calcined Clay (yellow). (**B**) The combined relative abundance of the top 20 ASVs commonly enriched in all soil treatments over time. Each point represents an individual sample. Patterns of enrichment were similar for both selection line (filled circles) and sterile-inoculated (open circles) microbiomes.

**Supplemental Figure 7.**
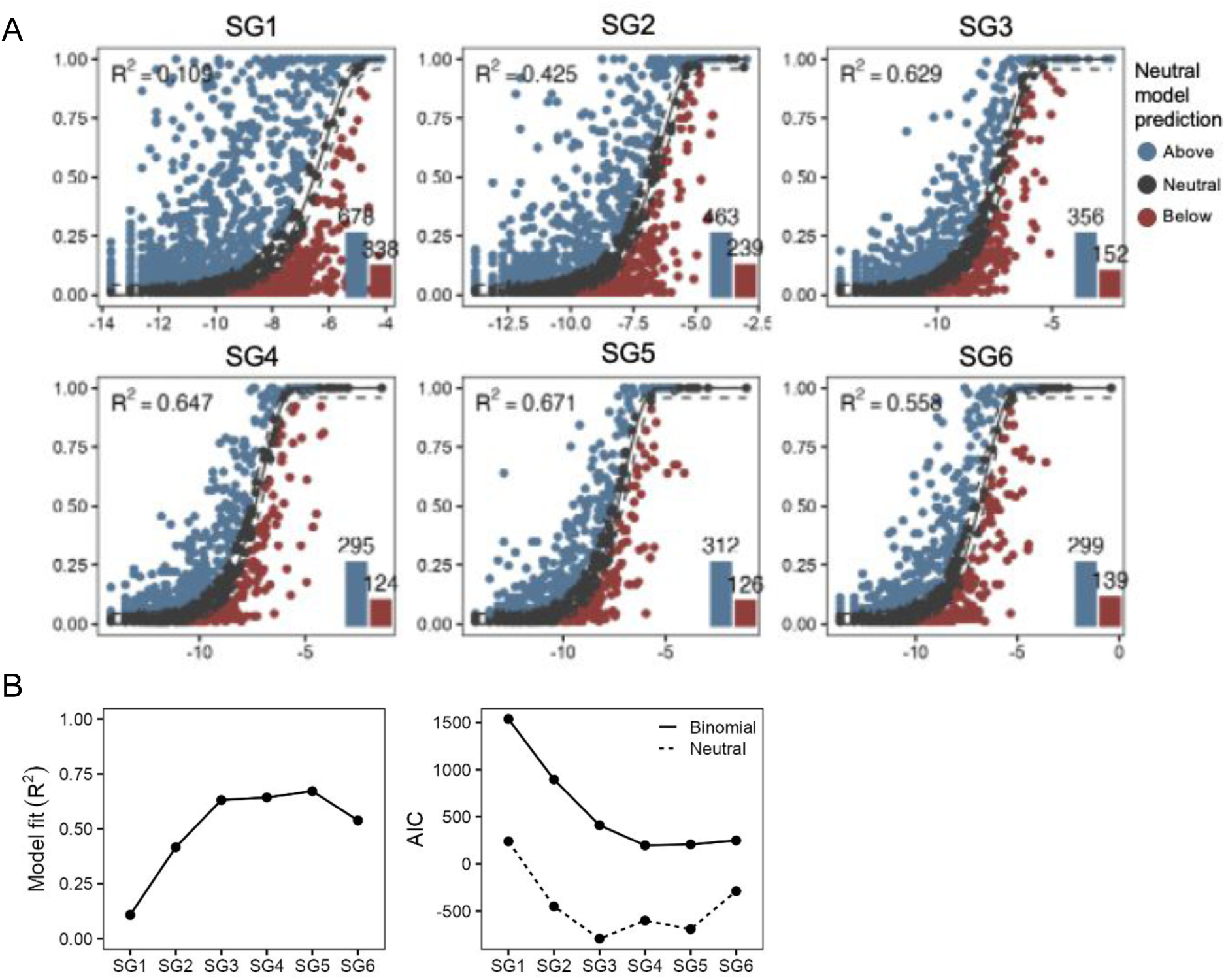
Neutral community models for Rice Field selection line microbiomes at each selection generation. (**A**) We fit ASV relative abundance data to neutral models using the previous generation’s community to parameterize migration and replacement rates. Gray points fall within the 95% confidence interval for abundance-occupancy values (x and y axes, respectively) expected if the community were assembled via neutral processes alone. Blue and red points indicate ASVs outside this prediction and therefore more likely experiencing positive and purifying selection, respectively. R-squared values for each plot indicate the overall model fit; blue and red bars in the bottom right corner of each plot show the total number of ASVs above and below neutral model predictions. (**B**) Given the improved fit of neutral models in later selection generations, neutral processes appear to become more important to microbiome assembly over time. Additionally, neutral models consistently outperformed binomial ones, which produced similar abundance-occupancy curves but without parameterization.

**Supplemental Figure 8.**
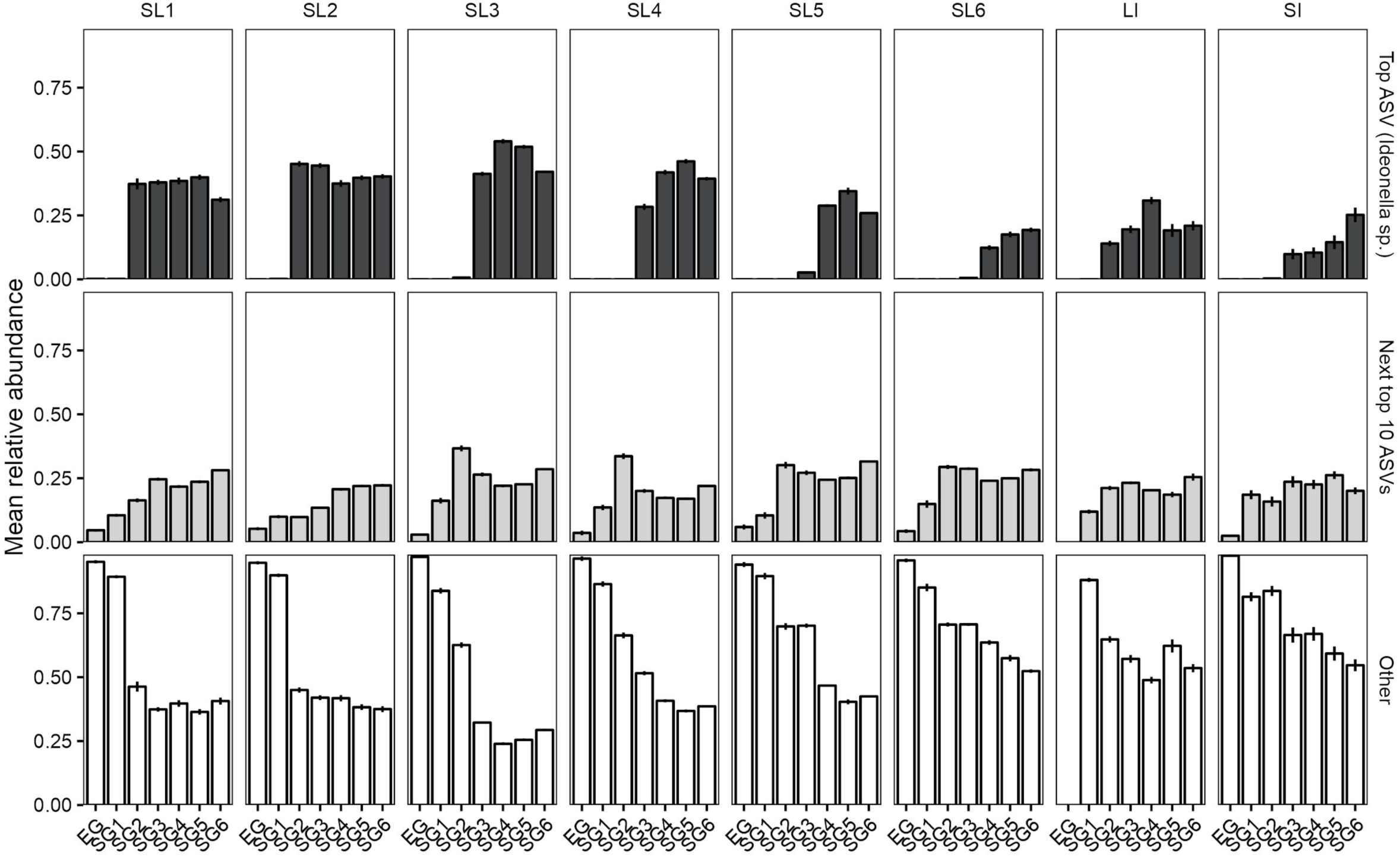
Most abundant ASVs within Rice Field microbiomes. One ASV in particular—identified as belonging to the genus *Ideonella—*rose to high abundance in Rice Field microbiomes, sometimes accounting for more than 50% of the relative abundance of the community. Often, the combined relative abundance of next top ten most abundant taxa was still less than that of *Ideonella* sp. Regardless, both showed patterns of enrichment over time in all selection lines as well as sterile- and live-inoculated microbiomes. Bars represent mean relative abundance +/- one standard error.

**Supplemental Figure 9.**
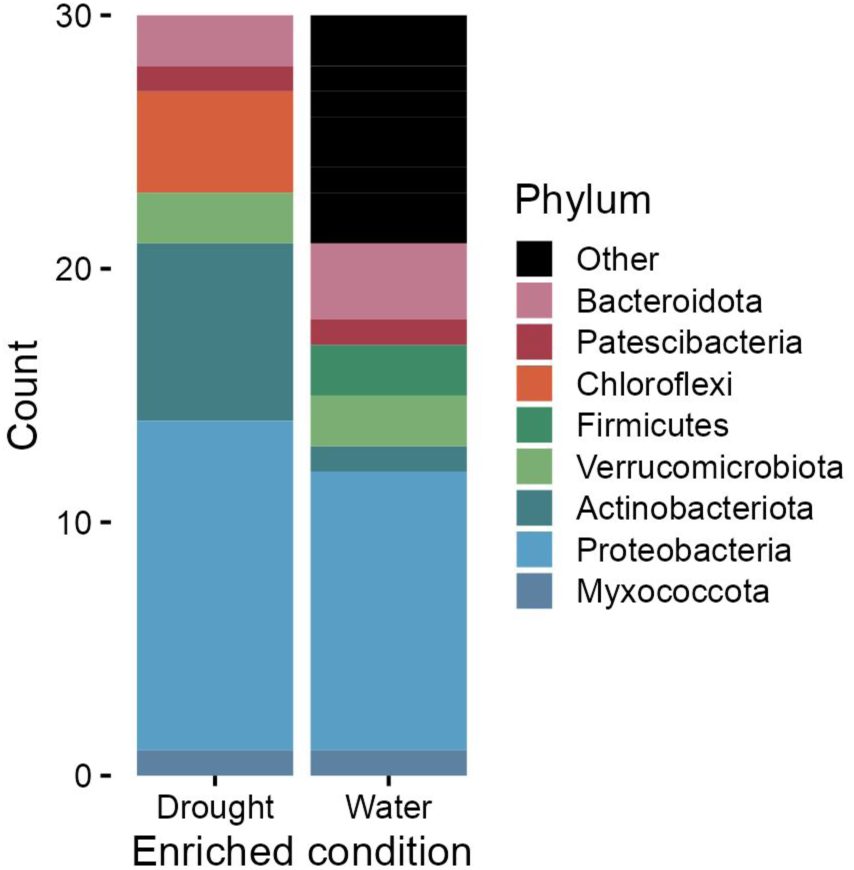
ASVs identified as significantly and consistently enriched in either selection line (drought) or live-inoculated (water) microbiomes. After filtering for FDR-corrected significant enrichment and an effect size greater than 0.5 (indicating significant and strong enrichment signal) we further subset to ASVs meeting these criteria in at least three selection generations. In total, 30 drought and 30 water indicators remained.

**Supplemental Figure 10.**
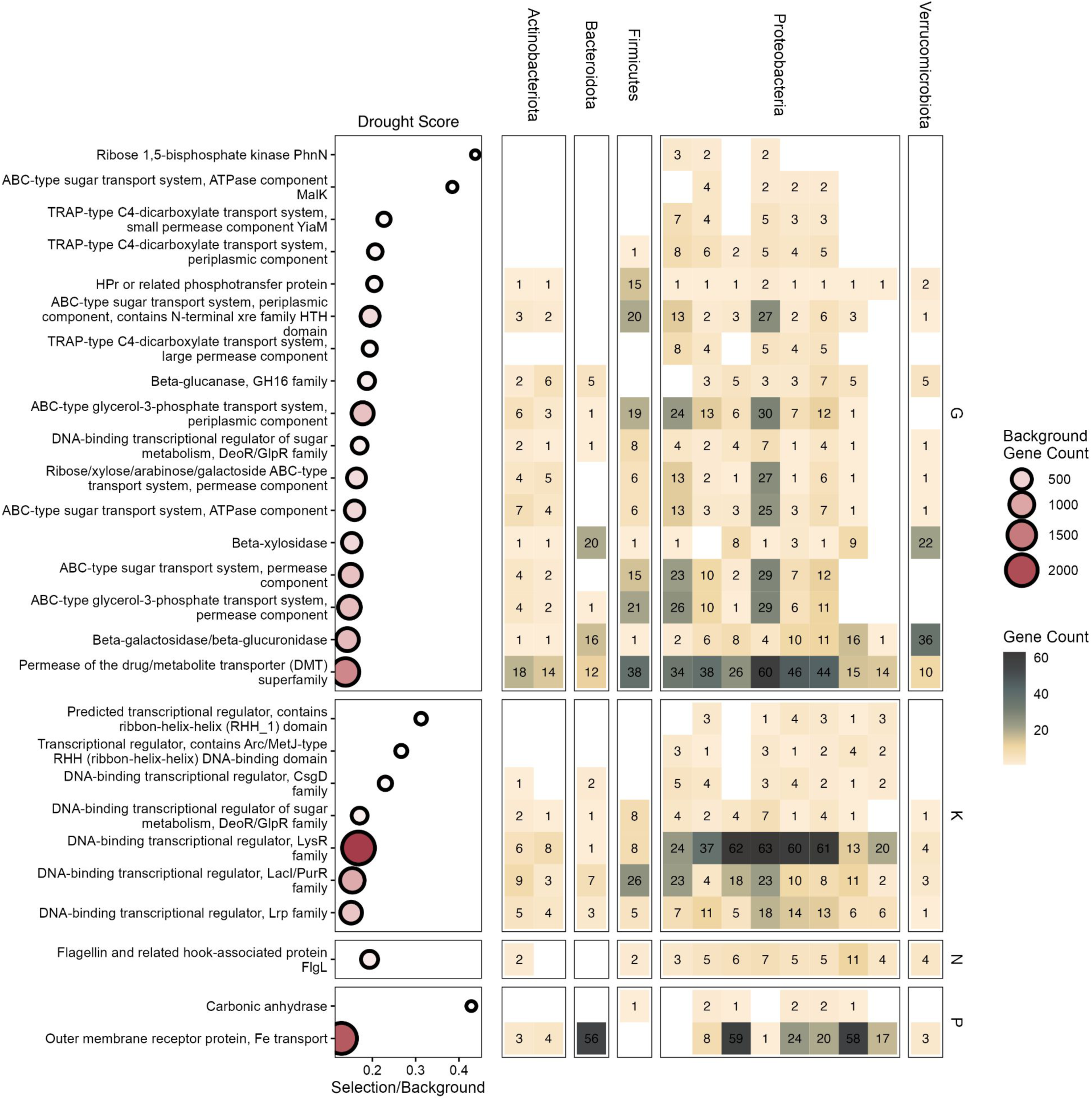
Enrichments of individual COGs within significantly enriched categories associated with drought score balance numerator taxa. As in Figure 7, Selection/Background refers to the ratio of gene counts among phenotype-associated taxa relative to all taxa, but for individual COGs rather than COG functional categories. All COGs shown were significantly enriched in balance taxa, even after applying multiple testing corrections scaled by the total number of COGs within each functional category. Each column in the heatmap represents an individual MAG; columns are grouped by phylum. Numbers within heatmap cells indicate the COG gene count for each MAG.

**Supplemental Figure 11.**
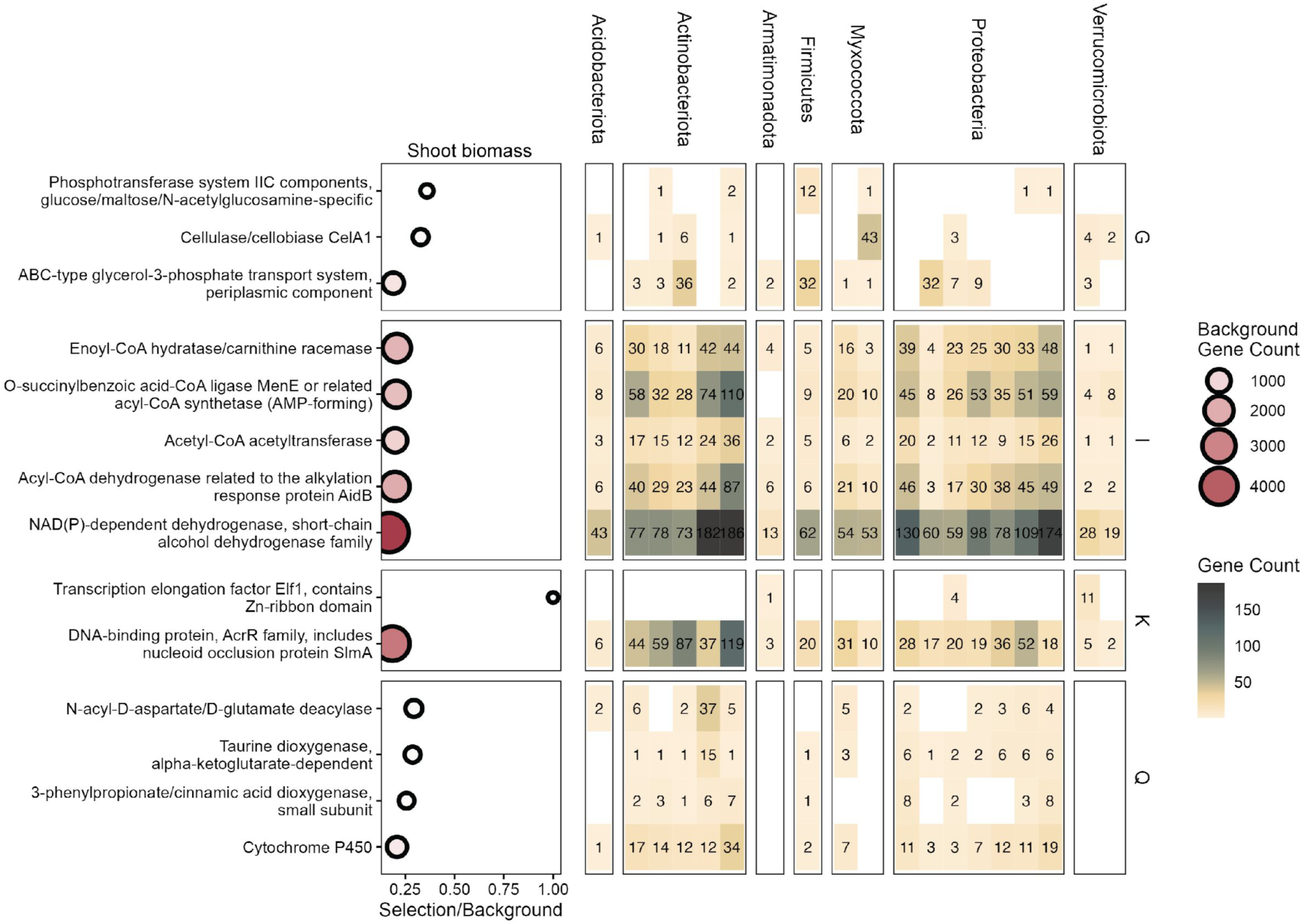
Enrichments of individual COGs within significantly enriched categories associated with shoot biomass balance numerator taxa. As in Figure 7, Selection/Background refers to the ratio of gene counts among phenotype-associated taxa relative to all taxa, but for individual COGs rather than COG functional categories. All COGs shown were significantly enriched in balance taxa, even after applying multiple testing corrections scaled by the total number of COGs within each functional category. Each column in the heatmap represents an individual MAG; columns are grouped by phylum. Numbers within heatmap cells indicate the COG gene count for each MAG.

**Supplemental Figure 12.**
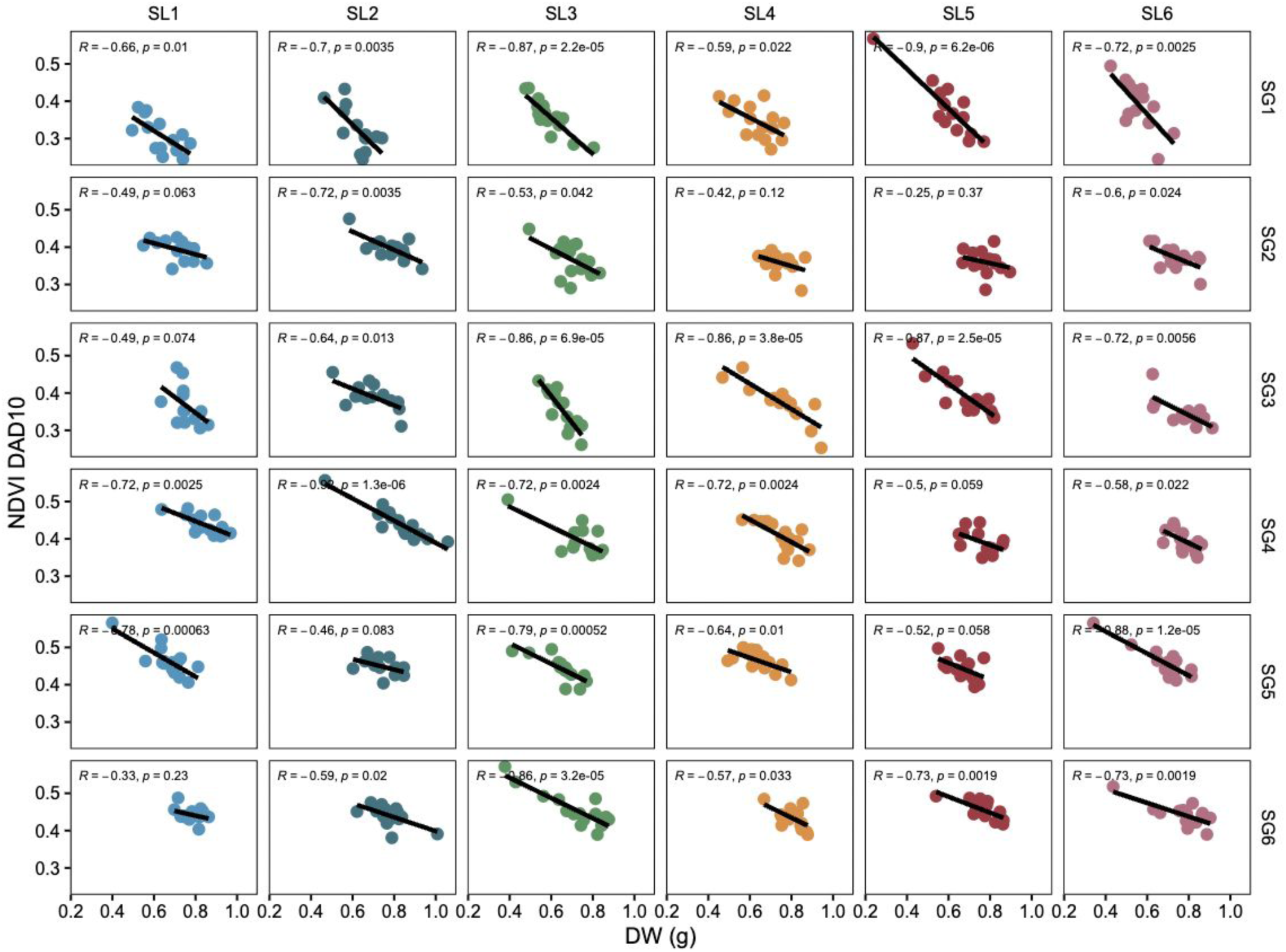
NDVI values for plants at 10 days after drought (DAD 10) decrease as rice shoot dry weight biomass increases. Data shown here are for Rice Field selection line plants only, but this pattern held true for other soil treatments and SI plants as well.

## References

1. Adamberg, K., and S. Adamberg. 2018. “Selection of Fast and Slow Growing Bacteria from Fecal Microbiota Using Continuous Culture with Changing Dilution Rate.” Microbial Ecology in Health and Disease 29 (1): 1549922. 10.1080/16512235.2018.1549922.

2. Albareda, Marta, Dulce Nombre Rodríguez-Navarro, and Francisco J. Temprano. 2009. “Soybean Inoculation: Dose, N Fertilizer Supplementation and Rhizobia Persistence in Soil.” Field Crops Research 113 (3): 352–56. 10.1016/j.fcr.2009.05.013.

3. Alneberg, Johannes, Brynjar Smári Bjarnason, Ino de Bruijn, Melanie Schirmer, Joshua Quick, Umer Z. Ijaz, Leo Lahti, Nicholas J. Loman, Anders F. Andersson, and Christopher Quince. 2014. “Binning Metagenomic Contigs by Coverage and Composition.” Nature Methods 11 : 1144–46. 10.1038/nmeth.3103.

4. Anderson, Marti J., Thomas O. Crist, Jonathan M. Chase, Mark Vellend, Brian D. Inouye, Amy L. Freestone, Nathan J. Sanders, et al. 2011. “Navigating the Multiple Meanings of β Diversity: A Roadmap for the Practicing Ecologist.” Ecology Letters 14 (1): 19–28. 10.1111/j.1461-0248.2010.01552.x.

5. Aslam, Ana, Zahir Ahmad Zahir, Hafiz Naeem Asghar, and Muhammad Shahid. 2018. “Effect of Carbonic Anhydrase-Containing Endophytic Bacteria on Growth and Physiological Attributes of Wheat under Water-Deficit Conditions.” Plant Production Science 21 (3): 244–55. 10.1080/1343943X.2018.1465348.

6. Bååth, Erland. 1996. “Adaptation of Soil Bacterial Communities to Prevailing pH in Different Soils.” FEMS Microbiology Ecology 19 (4): 227–37. 10.1111/j.1574-6941.1996.tb00215.x.

7. Bell, Thomas. 2010. “Experimental Tests of the Bacterial Distance-Decay Relationship.” The ISME Journal 4 (11): 1357–65. 10.1038/ismej.2010.77.

8. Belotte, Dorothée, Jean-Baptiste Curien, R. Craig Maclean, and Graham Bell. 2003. “An Experimental Test of Local Adaptation in Soil Bacteria.” Evolution; International Journal of Organic Evolution 57 (1): 27–36. 10.1111/j.0014-3820.2003.tb00213.x.

9. Bilal, Lubna, Sajjad Asaf, Muhammad Hamayun, Humaira Gul, Amjad Iqbal, Ihsan Ullah, In-Jung Lee, and Anwar Hussain. 2018. “Plant Growth Promoting Endophytic Fungi Asprgillus Fumigatus TS1 and Fusarium Proliferatum BRL1 Produce Gibberellins and Regulates Plant Endogenous Hormones.” Symbiosis 76 (2): 117–27. 10.1007/s13199-018-0545-4.

10. Bokulich, Nicholas A., Benjamin D. Kaehler, Jai Ram Rideout, Matthew Dillon, Evan Bolyen, Rob Knight, Gavin A. Huttley, and J. Gregory Caporaso. 2018. “Optimizing Taxonomic Classification of Marker-Gene Amplicon Sequences with QIIME 2’s q2-Feature-Classifier Plugin.” Microbiome 6 (1): 90. 10.1186/s40168-018-0470-z.

11. Bokulich, Nicholas A., John H. Thorngate, Paul M. Richardson, and David A. Mills. 2014. “Microbial Biogeography of Wine Grapes Is Conditioned by Cultivar, Vintage, and Climate.” Proceedings of the National Academy of Sciences of the United States of America 111 (1): E139–48. 10.1073/pnas.1317377110.

12. Bolyen, Evan, Jai Ram Rideout, Matthew R. Dillon, Nicholas A. Bokulich, Christian C. Abnet, Gabriel A. Al-Ghalith, Harriet Alexander, et al. 2019. “Reproducible, Interactive, Scalable and Extensible Microbiome Data Science Using QIIME 2.” Nature Biotechnology 37 (8): 852–57. 10.1038/s41587-019-0209-9.

13. Bradáčová, Klára, Andrea S. Florea, Asher Bar-Tal, Dror Minz, Uri Yermiyahu, Raneen Shawahna, Judith Kraut-Cohen, et al. 2019. “Microbial Consortia versus Single-Strain Inoculants: An Advantage in PGPM-Assisted Tomato Production?” Agronomy 9 (2): 105. 10.3390/agronomy9020105.

14. Burns, Adam R., W. Zac Stephens, Keaton Stagaman, Sandi Wong, John F. Rawls, Karen Guillemin, and Brendan Jm Bohannan. 2016. “Contribution of Neutral Processes to the Assembly of Gut Microbial Communities in the Zebrafish over Host Development.” The ISME Journal 10 (3): 655–64. 10.1038/ismej.2015.142.

15. Callahan, Benjamin J., Paul J. McMurdie, Michael J. Rosen, Andrew W. Han, Amy Jo A. Johnson, and Susan P. Holmes. 2016. “DADA2: High-Resolution Sample Inference from Illumina Amplicon Data.” Nature Methods 13 (7): 581–83. 10.1038/nmeth.3869.

16. Cantalapiedra, Carlos P., Ana Hernández-Plaza, Ivica Letunic, Peer Bork, and Jaime Huerta-Cepas. 2021. “eggNOG-Mapper v2: Functional Annotation, Orthology Assignments, and Domain Prediction at the Metagenomic Scale.” Molecular Biology and Evolution 38 : 5825–29. 10.1093/molbev/msab293.

17. Chaumeil, Pierre-Alain, Aaron J. Mussig, Philip Hugenholtz, and Donovan H. Parks. 2022. “GTDB-Tk v2: Memory Friendly Classification with the Genome Taxonomy Database.” Bioinformatics 38 (23): 5315–16. 10.1093/bioinformatics/btac672.

18. Chhetri, Geeta, Inhyup Kim, Minchung Kang, Jiyoun Kim, Yoonseop So, and Taegun Seo. 2022. “Devosia Rhizoryzae Sp. Nov., and Devosia Oryziradicis Sp. Nov., Novel Plant Growth Promoting Members of the Genus Devosia, Isolated from the Rhizosphere of Rice Plants.” Journal of Microbiology 60 (1): 1–10. 10.1007/s12275-022-1474-8.

19. Chhetri, Geeta, Inhyup Kim, Minchung Kang, Yoonseop So, Jiyoun Kim, and Taegun Seo. 2022. “An Isolated Arthrobacter Sp. Enhances Rice (Oryza Sativa L.) Plant Growth.” Microorganisms 10 (6). 10.3390/microorganisms10061187.

20. Debray, Reena, Robin A. Herbert, Alexander L. Jaffe, Alexander Crits-Christoph, Mary E. Power, and Britt Koskella. 2022. “Priority Effects in Microbiome Assembly.” Nature Reviews. Microbiology 20 (2): 109–21. 10.1038/s41579-021-00604-w.

21. Delgado-Baquerizo, Manuel, Fernando T. Maestre, Peter B. Reich, Thomas C. Jeffries, Juan J. Gaitan, Daniel Encinar, Miguel Berdugo, Colin D. Campbell, and Brajesh K. Singh. 2016. “Microbial Diversity Drives Multifunctionality in Terrestrial Ecosystems.” Nature Communications 7 (January): 10541. 10.1038/ncomms10541.

22. De Sousa Fabiano Gama, Mielke Kamila Cabral, Marques Caldeira Dany Roberta, Divan Baldani Vera Lucia, Baldani Jose Ivo, da Silva Roberlan Ferreira, Balbinot Ernando, and Klein Vanderley Antonio Chorobura. 2022. “Genetic Diversity and Inoculation of Plant-Growth Promoting Diazotrophic Bacteria for Production of ‘Eucalyptus Urophylla’ Seedlings.” Australian Journal of Crop Science 16 (1): 35–44. 10.3316/informit.643092702441833.

23. Díaz-García, Laura, Sixing Huang, Cathrin Spröer, Rocío Sierra-Ramírez, Boyke Bunk, Jörg Overmann, and Diego Javier Jiménez. 2021. “Dilution-to-Stimulation/Extinction Method: A Combination Enrichment Strategy To Develop a Minimal and Versatile Lignocellulolytic Bacterial Consortium.” Applied and Environmental Microbiology 87 (2). 10.1128/AEM.02427-20.

24. Durrant, W. E., and X. Dong. 2004. “Systemic Acquired Resistance.” Annual Review of Phytopathology 42: 185–209. 10.1146/annurev.phyto.42.040803.140421.

25. Evans, Sarah, Jennifer B. H. Martiny, and Steven D. Allison. 2017. “Effects of Dispersal and Selection on Stochastic Assembly in Microbial Communities.” The ISME Journal 11 (1): 176–85. 10.1038/ismej.2016.96.

26. Fahlgren, Noah, Maximilian Feldman, Malia A. Gehan, Melinda S. Wilson, Christine Shyu, Douglas W. Bryant, Steven T. Hill, et al. 2015. “A Versatile Phenotyping System and Analytics Platform Reveals Diverse Temporal Responses to Water Availability in Setaria.” Molecular Plant 8 (10): 1520–35. 10.1016/j.molp.2015.06.005.

27. Fan, Di, and Donald L. Smith. 2022. “Mucilaginibacter Sp. K Improves Growth and Induces Salt Tolerance in Nonhost Plants via Multilevel Mechanisms.” Frontiers in Plant Science 13 (June): 938697. 10.3389/fpls.2022.938697.

28. Faust, Karoline, and Jeroen Raes. 2012. “Microbial Interactions: From Networks to Models.” Nature Reviews. Microbiology 10 (8): 538–50. 10.1038/nrmicro2832.

29. Fierer, Noah, and Robert B. Jackson. 2006. “The Diversity and Biogeography of Soil Bacterial Communities.” Proceedings of the National Academy of Sciences of the United States of America 103 (3): 626–31. 10.1073/pnas.0507535103.

30. Finkel, Omri M., Gabriel Castrillo, Sur Herrera Paredes, Isai Salas González, and Jeffery L. Dangl. 2017. “Understanding and Exploiting Plant Beneficial Microbes.” Current Opinion in Plant Biology 38 (August): 155–63. 10.1016/j.pbi.2017.04.018.

31. Fitzpatrick, Connor R., Julia Copeland, Pauline W. Wang, David S. Guttman, Peter M. Kotanen, and Marc T. J. Johnson. 2018. “Assembly and Ecological Function of the Root Microbiome across Angiosperm Plant Species.” Proceedings of the National Academy of Sciences of the United States of America 115 (6): E1157–65. 10.1073/pnas.1717617115.

32. Fodelianakis, Stilianos, Alexander Lorz, Adriana Valenzuela-Cuevas, Alan Barozzi, Jenny Marie Booth, and Daniele Daffonchio. 2019. “Dispersal Homogenizes Communities via Immigration Even at Low Rates in a Simplified Synthetic Bacterial Metacommunity.” Nature Communications 10 (1): 1314. 10.1038/s41467-019-09306-7.

33. Franche, Claudine, Kristina Lindström, and Claudine Elmerich. 2009. “Nitrogen-Fixing Bacteria Associated with Leguminous and Non-Leguminous Plants.” Plant and Soil 321 (1): 35–59. 10.1007/s11104-008-9833-8.

34. Frantz, Jonathan M., and Bruce Bugbee. 2002. “Anaerobic Conditions Improve Germination of a Gibberellic Acid Deficient Rice.” Crop Science 42 (2): 651–54. 10.2135/cropsci2002.6510.

35. Friedman, Jonathan, Logan M. Higgins, and Jeff Gore. 2017. “Community Structure Follows Simple Assembly Rules in Microbial Microcosms.” Nature Ecology & Evolution 1 (5): 109. 10.1038/s41559-017-0109.

36. Friesen, Maren L., Stephanie S. Porter, Scott C. Stark, Eric J. von Wettberg, Joel L. Sachs, and Esperanza Martinez-Romero. 2011. “Microbially Mediated Plant Functional Traits.” Annual Review of Ecology, Evolution, and Systematics 42 (1): 23–46. 10.1146/annurev-ecolsys-102710-145039.

37. Fukami, Josiane, Paula Cerezini, and Mariangela Hungria. 2018. “Azospirillum: Benefits That Go Far beyond Biological Nitrogen Fixation.” AMB Express 8 (1): 73. 10.1186/s13568-018-0608-1.

38. Galperin, Michael Y., Yuri I. Wolf, Kira S. Makarova, Roberto Vera Alvarez, David Landsman, and Eugene V. Koonin. 2021. “COG Database Update: Focus on Microbial Diversity, Model Organisms, and Widespread Pathogens.” Nucleic Acids Research 49 (D1): D274–81. 10.1093/nar/gkaa1018.

39. Garcia, Joshua, Maria Gannett, Liping Wei, Liang Cheng, Shengyuan Hu, Jed Sparks, James Giovannoni, and Jenny Kao-Kniffin. 2022. “Selection Pressure on the Rhizosphere Microbiome Can Alter Nitrogen Use Efficiency and Seed Yield in Brassica Rapa.” Communications Biology 5 (1): 959. 10.1038/s42003-022-03860-5.

40. Glick, Bernard R. 2005. “Modulation of Plant Ethylene Levels by the Bacterial Enzyme ACC Deaminase.” FEMS Microbiology Letters 251 (1): 1–7. 10.1016/j.femsle.2005.07.030.

41. Herrera Paredes, Sur, Tianxiang Gao, Theresa F. Law, Omri M. Finkel, Tatiana Mucyn, Paulo José Pereira Lima Teixeira, Isaí Salas González, et al. 2018. “Design of Synthetic Bacterial Communities for Predictable Plant Phenotypes.” PLoS Biology 16 (2): e2003962. 10.1371/journal.pbio.2003962.

42. Huang, Sha, Lina Tang, Joseph P. Hupy, Yang Wang, and Guofan Shao. 2021. “A Commentary Review on the Use of Normalized Difference Vegetation Index (NDVI) in the Era of Popular Remote Sensing.” Research Journal of Forestry 32 (1): 1–6. 10.1007/s11676-020-01155-1.

43. Johnson, Nancy Collins, Gail W. T. Wilson, Matthew A. Bowker, Jacqueline A. Wilson, and R. Michael Miller. 2010. “Resource Limitation Is a Driver of Local Adaptation in Mycorrhizal Symbioses.” Proceedings of the National Academy of Sciences of the United States of America 107 (5): 2093–98. 10.1073/pnas.0906710107.

44. Jones, Eric W., and Jean M. Carlson. 2019. “Steady-State Reduction of Generalized Lotka-Volterra Systems in the Microbiome.” Physical Review. E 99 (3-1): 032403. 10.1103/PhysRevE.99.032403.

45. Kalachova, Tetiana, Barbora Jindřichová, Lenka Burketová, Cécile Monard, Manuel Blouin, Samuel Jacquiod, Eric Ruelland, and Ruben Puga-Freitas. 2022. “Controlled Natural Selection of Soil Microbiome through Plant-Soil Feedback Confers Resistance to a Foliar Pathogen.” *Plant and Soil*, July. 10.1007/s11104-022-05597-w.

46. Kang, Dongwan D., Feng Li, Edward Kirton, Ashleigh Thomas, Rob Egan, Hong An, and Zhong Wang. 2019. “MetaBAT 2: An Adaptive Binning Algorithm for Robust and Efficient Genome Reconstruction from Metagenome Assemblies.” PeerJ 7 (July): e7359. 10.7717/peerj.7359.

47. Karmakar, Joydip, Sayani Goswami, Krishnendu Pramanik, Tushar Kanti Maiti, Rup Kumar Kar, and Narottam Dey. 2021. “Growth Promoting Properties of Mycobacterium and Bacillus on Rice Plants under Induced Drought.” Plant Science Today 8 (1): 49–57. 10.14719/pst.2021.8.1.965.

48. Kaur, Simranjit, Eleonora Egidi, Zhiguang Qiu, Catriona A. Macdonald, Jay Prakash Verma, Pankaj Trivedi, Juntao Wang, Hongwei Liu, and Brajesh K. Singh. 2022. “Synthetic Community Improves Crop Performance and Alters Rhizosphere Microbial Communities.” Journal of Sustainable Agriculture and Environment 1 (2): 118–31. 10.1002/sae2.12017.

49. King, William L., Laura M. Kaminsky, Maria Gannett, Grant L. Thompson, Jenny Kao-Kniffin, and Terrence H. Bell. 2022. “Soil Salinization Accelerates Microbiome Stabilization in Iterative Selections for Plant Performance.” The New Phytologist 234 (6): 2101–10. 10.1111/nph.17774.

50. Korotkevich, Gennady, Vladimir Sukhov, Nikolay Budin, Boris Shpak, Maxim N. Artyomov, and Alexey Sergushichev. 2016. “Fast Gene Set Enrichment Analysis.” bioRxiv. bioRxiv. 10.1101/060012.

51. Kraemer, Susanne A., and Primrose J. Boynton. 2017. “Evidence for Microbial Local Adaptation in Nature.” Molecular Ecology 26 (7): 1860–76. 10.1111/mec.13958.

52. Kraemer, Susanne A., and Rees Kassen. 2015. “Patterns of Local Adaptation in Space and Time among Soil Bacteria.” The American Naturalist 185 (3): 317–31. 10.1086/679585.

53. Kramer, Jos, Özhan Özkaya, and Rolf Kümmerli. 2020. “Bacterial Siderophores in Community and Host Interactions.” Nature Reviews. Microbiology 18 (3): 152–63. 10.1038/s41579-019-0284-4.

54. Kumar, Akhilesh, Saurabh Singh, Anand Kumar Gaurav, Sudhakar Srivastava, and Jay Prakash Verma. 2020. “Plant Growth-Promoting Bacteria: Biological Tools for the Mitigation of Salinity Stress in Plants.” Frontiers in Microbiology 11 (July): 1216. 10.3389/fmicb.2020.01216.

55. Kuzyakov, Yakov, and Bahar S. Razavi. 2019. “Rhizosphere Size and Shape: Temporal Dynamics and Spatial Stationarity.” Soil Biology & Biochemistry 135 (August): 343–60. 10.1016/j.soilbio.2019.05.011.

56. Lee, Michael D. 2019. “GToTree: A User-Friendly Workflow for Phylogenomics.” Bioinformatics 35 (20): 4162–64. 10.1093/bioinformatics/btz188.

57. Li, Dinghua, Chi-Man Liu, Ruibang Luo, Kunihiko Sadakane, and Tak-Wah Lam. 2015. “MEGAHIT: An Ultra-Fast Single-Node Solution for Large and Complex Metagenomics Assembly via Succinct de Bruijn Graph.” Bioinformatics 31 (10): 1674–76. 10.1093/bioinformatics/btv033.

58. Li, Dinghua, Ruibang Luo, Chi-Man Liu, Chi-Ming Leung, Hing-Fung Ting, Kunihiko Sadakane, Hiroshi Yamashita, and Tak-Wah Lam. 2016. “MEGAHIT v1.0: A Fast and Scalable Metagenome Assembler Driven by Advanced Methodologies and Community Practices.” Methods 102 (June): 3–11. 10.1016/j.ymeth.2016.02.020.

59. Li, Heng. 2013. “Aligning Sequence Reads, Clone Sequences and Assembly Contigs with BWA-MEM.” *arXiv [q-bio.GN]*. arXiv. http://arxiv.org/abs/1303.3997.

60. Liu, Hongwei, Laura E. Brettell, Zhiguang Qiu, and Brajesh K. Singh. 2020. “Microbiome-Mediated Stress Resistance in Plants.” Trends in Plant Science 25 (8): 733–43. 10.1016/j.tplants.2020.03.014.

61. Liu, Hongwei, Zhiguang Qiu, Jun Ye, Jay Prakash Verma, Jiayu Li, and Brajesh K. Singh. 2022. “Effective Colonisation by a Bacterial Synthetic Community Promotes Plant Growth and Alters Soil Microbial Community.” Journal of Sustainable Agriculture and Environment 1 (1): 30–42. 10.1002/sae2.12008.

62. López, José L., Arista Fourie, Sanne W. M. Poppeliers, Nikolaos Pappas, Juan J. Sánchez-Gil, Ronnie de Jonge, and Bas E. Dutilh. 2023. “Growth Rate Is a Dominant Factor Predicting the Rhizosphere Effect.” The ISME Journal 17 (9): 1396–1405. 10.1038/s41396-023-01453-6.

63. Martin, Marcel. 2011. “Cutadapt Removes Adapter Sequences from High-Throughput Sequencing Reads.” EMBnet.journal 17 (1): 10–12. 10.14806/ej.17.1.200.

64. Martino, Cameron, James T. Morton, Clarisse A. Marotz, Luke R. Thompson, Anupriya Tripathi, Rob Knight, and Karsten Zengler. 2019. “A Novel Sparse Compositional Technique Reveals Microbial Perturbations.” mSystems 4 (1). 10.1128/mSystems.00016-19.

65. McCarty, Nicholas S., and Rodrigo Ledesma-Amaro. 2019. “Synthetic Biology Tools to Engineer Microbial Communities for Biotechnology.” Trends in Biotechnology 37 (2): 181–97. 10.1016/j.tibtech.2018.11.002.

66. Morton, James T., Jon Sanders, Robert A. Quinn, Daniel McDonald, Antonio Gonzalez, Yoshiki Vázquez-Baeza, Jose A. Navas-Molina, et al. 2017. “Balance Trees Reveal Microbial Niche Differentiation.” mSystems 2 (1). 10.1128/mSystems.00162-16.

67. Mueller, U. G., and J. L. Sachs. 2015. “Engineering Microbiomes to Improve Plant and Animal Health.” Trends in Microbiology 23 (10): 606–17. 10.1016/j.tim.2015.07.009.

68. Mueller, Ulrich G., and Timothy A. Linksvayer. 2022. “Microbiome Breeding: Conceptual and Practical Issues.” *Trends in Microbiology*, May. 10.1016/j.tim.2022.04.003.

69. Naylor, Dan, Stephanie DeGraaf, Elizabeth Purdom, and Devin Coleman-Derr. 2017. “Drought and Host Selection Influence Bacterial Community Dynamics in the Grass Root Microbiome.” The ISME Journal 11 (12): 2691–2704. 10.1038/ismej.2017.118.

70. Niu, Ben, Joseph Nathaniel Paulson, Xiaoqi Zheng, and Roberto Kolter. 2017. “Simplified and Representative Bacterial Community of Maize Roots.” Proceedings of the National Academy of Sciences of the United States of America 114 (12): E2450–59. 10.1073/pnas.1616148114.

71. Niu, Shuqi, Yan Gao, Huixian Zi, Ying Liu, Xuanming Liu, Xianqiu Xiong, Qingqing Yao, et al. 2022. “The Osmolyte-Producing Endophyte Streptomyces Albidoflavus OsiLf-2 Induces Drought and Salt Tolerance in Rice via a Multi-Level Mechanism.” The Crop Journal 10 (2): 375–86. 10.1016/j.cj.2021.06.008.

72. Olm, Matthew R., Christopher T. Brown, Brandon Brooks, and Jillian F. Banfield. 2017. “dRep: A Tool for Fast and Accurate Genomic Comparisons That Enables Improved Genome Recovery from Metagenomes through de-Replication.” The ISME Journal 11 (12): 2864–68. 10.1038/ismej.2017.126.

73. Orozco-Mosqueda, Ma Del Carmen, Gustavo Santoyo, and Bernard R. Glick. 2023. “Recent Advances in the Bacterial Phytohormone Modulation of Plant Growth.” Plants 12 (3). 10.3390/plants12030606.

74. Panda, Debabrata, Swati Sakambari Mishra, and Prafulla Kumar Behera. 2021. “Drought Tolerance in Rice: Focus on Recent Mechanisms and Approaches.” Rice Science 28 (2): 119–32. 10.1016/j.rsci.2021.01.002.

75. Pandey, Veena, and Alok Shukla. 2015. “Acclimation and Tolerance Strategies of Rice under Drought Stress.” Rice Science 22 (4): 147–61. 10.1016/j.rsci.2015.04.001.

76. Panke-Buisse, Kevin, Angela C. Poole, Julia K. Goodrich, Ruth E. Ley, and Jenny Kao-Kniffin. 2015. “Selection on Soil Microbiomes Reveals Reproducible Impacts on Plant Function.” The ISME Journal 9 (4): 980–89. 10.1038/ismej.2014.196.

77. Pedregosa, Fabian, Gaël Varoquaux, Alexandre Gramfort, Vincent Michel, Bertrand Thirion, Olivier Grisel, Mathieu Blondel, et al. 2012. “Scikit-Learn: Machine Learning in Python.” *arXiv [cs.LG]*. arXiv. https://www.jmlr.org/papers/volume12/pedregosa11a/pedregosa11a.pdf.

78. Pieterse, Corné M. J., Christos Zamioudis, Roeland L. Berendsen, David M. Weller, Saskia C. M. Van Wees, and Peter A. H. M. Bakker. 2014. “Induced Systemic Resistance by Beneficial Microbes.” Annual Review of Phytopathology 52 (June): 347–75. 10.1146/annurev-phyto-082712-102340.

79. Qian, Haoyu, Xiangchen Zhu, Shan Huang, Bruce Linquist, Yakov Kuzyakov, Reiner Wassmann, Kazunori Minamikawa, et al. 2023. “Greenhouse Gas Emissions and Mitigation in Rice Agriculture.” Nature Reviews Earth & Environment 4 (10): 716–32. 10.1038/s43017-023-00482-1.

80. Quast, Christian, Elmar Pruesse, Pelin Yilmaz, Jan Gerken, Timmy Schweer, Pablo Yarza, Jörg Peplies, and Frank Oliver Glöckner. 2013. “The SILVA Ribosomal RNA Gene Database Project: Improved Data Processing and Web-Based Tools.” Nucleic Acids Research 41 (Database issue): D590–96. 10.1093/nar/gks1219.

81. Richardson, Alan E., and Richard J. Simpson. 2011. “Soil Microorganisms Mediating Phosphorus Availability Update on Microbial Phosphorus.” Plant Physiology 156 (3): 989–96. 10.1104/pp.111.175448.

82. Rivas, Raúl, Anne Willems, Nanjappa S. Subba-Rao, Pedro F. Mateos, Frank B. Dazzo, Reiner M. Kroppenstedt, Eustoquio Martínez-Molina, Monique Gillis, and Encarna Velázquez. 2003. “Description of Devosia Neptuniae Sp. Nov. That Nodulates and Fixes Nitrogen in Symbiosis with Neptunia Natans, an Aquatic Legume from India.” Systematic and Applied Microbiology 26 (1): 47–53. 10.1078/072320203322337308.

83. Rivera-Pinto, J., J. J. Egozcue, V. Pawlowsky-Glahn, R. Paredes, M. Noguera-Julian, and M. L. Calle. 2018. “Balances: A New Perspective for Microbiome Analysis.” mSystems 3 (4). 10.1128/mSystems.00053-18.

84. Rodrigues, Elisete Pains, Luciana Santos Rodrigues, André Luiz Martinez de Oliveira, Vera Lucia Divan Baldani, Kátia Regina dos Santos Teixeira, Segundo Urquiaga, and Veronica Massena Reis. 2008. “Azospirillum Amazonense Inoculation: Effects on Growth, Yield and N2 Fixation of Rice (Oryza Sativa L.).” Plant and Soil 302 (1): 249–61. 10.1007/s11104-007-9476-1.

85. Rodríguez, Rodrigo, Patricio J. Barra, Giovanni Larama, Víctor J. Carrion, María de la Luz Mora, Lauren Hale, and Paola Durán. 2023. “Microbiome Engineering Optimized by Antarctic Microbiota to Support a Plant Host under Water Deficit.” Frontiers in Plant Science 14 (September): 1241612. 10.3389/fpls.2023.1241612.

86. Rouse, John Wilson, Rüdiger H. Haas, John A. Schell, Donald W. Deering, and Others. 1974. “Monitoring Vegetation Systems in the Great Plains with ERTS.” NASA. Special Publication 351 (1): 309. https://books.google.com/books?hl=en&lr=&id=e00CAAAAIAAJ&oi=fnd&pg=PA309&ots=JTTzbUCrWe&sig=fDJBu6FeoRGhZHIYk9mOevxqpo0.

87. Ruíz-Sánchez, Michel, Elisabet Armada, Yaumara Muñoz, Inés E. García de Salamone, Ricardo Aroca, Juan Manuel Ruíz-Lozano, and Rosario Azcón. 2011. “Azospirillum and Arbuscular Mycorrhizal Colonization Enhance Rice Growth and Physiological Traits under Well-Watered and Drought Conditions.” Journal of Plant Physiology 168 (10): 1031–37. 10.1016/j.jplph.2010.12.019.

88. Russ, Dor, Connor R. Fitzpatrick, Paulo J. P. L. Teixeira, and Jeffery L. Dangl. 2023. “Deep Discovery Informs Difficult Deployment in Plant Microbiome Science.” Cell 186 (21): 4496–4513. 10.1016/j.cell.2023.08.035.

89. Sansinenea, Estibaliz. 2019. “Bacillus Spp.: As Plant Growth-Promoting Bacteria.” In Secondary Metabolites of Plant Growth Promoting Rhizomicroorganisms: Discovery and Applications, edited by Harikesh Bahadur Singh, Chetan Keswani, M. S. Reddy, Estibaliz Sansinenea, and Carlos García-Estrada, 225–37. Singapore: Springer Singapore. 10.1007/978-981-13-5862-3_11.

90. Santos-Medellín, Christian, Joseph Edwards, Zachary Liechty, Bao Nguyen, and Venkatesan Sundaresan. 2017. “Drought Stress Results in a Compartment-Specific Restructuring of the Rice Root-Associated Microbiomes.” mBio 8 (4). 10.1128/mBio.00764-17.

91. Schepers, J. S., T. M. Blackmer, W. W. Wilhelm, and M. Resende. 1996. “Transmittance and Reflectance Measurements of CornLeaves from Plants with Different Nitrogen and Water Supply.” Journal of Plant Physiology 148 (5): 523–29. 10.1016/S0176-1617(96)80071-X.

92. Schlatter, Daniel, Linda Kinkel, Linda Thomashow, David Weller, and Timothy Paulitz. 2017. “Disease Suppressive Soils: New Insights from the Soil Microbiome.” Phytopathology 107 (11): 1284–97. 10.1094/PHYTO-03-17-0111-RVW.

93. Seemann, Torsten. 2014. “Prokka: Rapid Prokaryotic Genome Annotation.” Bioinformatics 30 (14): 2068–69. 10.1093/bioinformatics/btu153.

94. Selim, Samy, Yasser M. Hassan, Ahmed M. Saleh, Talaat H. Habeeb, and Hamada AbdElgawad. 2019. “Actinobacterium Isolated from a Semi-Arid Environment Improves the Drought Tolerance in Maize (Zea Mays L.).” Plant Physiology and Biochemistry: PPB / Societe Francaise de Physiologie Vegetale 142 (September): 15–21. 10.1016/j.plaphy.2019.06.029.

95. Sessitsch, Angela, Nikolaus Pfaffenbichler, and Birgit Mitter. 2019. “Microbiome Applications from Lab to Field: Facing Complexity.” Trends in Plant Science 24 (3): 194–98. 10.1016/j.tplants.2018.12.004.

96. Shayanthan, Ambihai, Patricia Ann C. Ordoñez, and Ivan John Oresnik. 2022. “The Role of Synthetic Microbial Communities (SynCom) in Sustainable Agriculture.” Frontiers in Agronomy 4. 10.3389/fagro.2022.896307.

97. Sieber, Christian M. K., Alexander J. Probst, Allison Sharrar, Brian C. Thomas, Matthias Hess, Susannah G. Tringe, and Jillian F. Banfield. 2018. “Recovery of Genomes from Metagenomes via a Dereplication, Aggregation and Scoring Strategy.” Nature Microbiology 3 (7): 836–43. 10.1038/s41564-018-0171-1.

98. Siebers, Meike, Mathias Brands, Vera Wewer, Yanjiao Duan, Georg Hölzl, and Peter Dörmann. 2016. “Lipids in Plant-Microbe Interactions.” Biochimica et Biophysica Acta 1861 (9 Pt B): 1379–95. 10.1016/j.bbalip.2016.02.021.

99. Sloan, William T., Mary Lunn, Stephen Woodcock, Ian M. Head, Sean Nee, and Thomas P. Curtis. 2006. “Quantifying the Roles of Immigration and Chance in Shaping Prokaryote Community Structure.” Environmental Microbiology 8 (4): 732–40. 10.1111/j.1462-2920.2005.00956.x.

100. Spaepen, Stijn. 2015. “Plant Hormones Produced by Microbes.” In Principles of Plant-Microbe Interactions: Microbes for Sustainable Agriculture, edited by Ben Lugtenberg, 247–56. Cham: Springer International Publishing. 10.1007/978-3-319-08575-3_26.

101. Spaepen, Stijn, and Jos Vanderleyden. 2011. “Auxin and Plant-Microbe Interactions.” Cold Spring Harbor Perspectives in Biology 3 (4). 10.1101/cshperspect.a001438.

102. Ustin, Susan L., and Stéphane Jacquemoud. 2020. “How the Optical Properties of Leaves Modify the Absorption and Scattering of Energy and Enhance Leaf Functionality.” In Remote Sensing of Plant Biodiversity, edited by Jeannine Cavender-Bares, John A. Gamon, and Philip A. Townsend, 349–84. Cham: Springer International Publishing. 10.1007/978-3-030-33157-3_14.

103. Vorholt, Julia A., Christine Vogel, Charlotte I. Carlström, and Daniel B. Müller. 2017. “Establishing Causality: Opportunities of Synthetic Communities for Plant Microbiome Research.” Cell Host & Microbe 22 (2): 142–55. 10.1016/j.chom.2017.07.004.

104. Wang, Boxi, and Shuichi Sugiyama. 2020. “Phylogenetic Signal of Host Plants in the Bacterial and Fungal Root Microbiomes of Cultivated Angiosperms.” The Plant Journal: For Cell and Molecular Biology 104 (2): 522–31. 10.1111/tpj.14943.

105. Wang, Honggui, Zhong Wei, Lijuan Mei, Jingxin Gu, Suisui Yin, Karoline Faust, Jeroen Raes, et al. 2017. “Combined Use of Network Inference Tools Identifies Ecologically Meaningful Bacterial Associations in a Paddy Soil.” Soil Biology & Biochemistry 105 (February): 227–35. 10.1016/j.soilbio.2016.11.029.

106. Weissman, J. L., Shengwei Hou, and Jed A. Fuhrman. 2021. “Estimating Maximal Microbial Growth Rates from Cultures, Metagenomes, and Single Cells via Codon Usage Patterns.” Proceedings of the National Academy of Sciences of the United States of America 118 (12). 10.1073/pnas.2016810118.

107. Whitham, Thomas G., Gerard J. Allan, Hillary F. Cooper, and Stephen M. Shuster. 2020. “Intraspecific Genetic Variation and Species Interactions Contribute to Community Evolution.” Annual Review of Ecology, Evolution, and Systematics 51 (1): 587–612. 10.1146/annurev-ecolsys-011720-123655.

108. Whitman, Thea, Rachel Neurath, Adele Perera, Ilexis Chu-Jacoby, Daliang Ning, Jizhong Zhou, Peter Nico, Jennifer Pett-Ridge, and Mary Firestone. 2018. “Microbial Community Assembly Differs across Minerals in a Rhizosphere Microcosm.” Environmental Microbiology 20 (12): 4444–60. 10.1111/1462-2920.14366.

109. Wright, Robyn J., Matthew I. Gibson, and Joseph A. Christie-Oleza. 2019. “Understanding Microbial Community Dynamics to Improve Optimal Microbiome Selection.” Microbiome 7 (1): 85. 10.1186/s40168-019-0702-x.

110. Wu, Yu-Wei, Blake A. Simmons, and Steven W. Singer. 2016. “MaxBin 2.0: An Automated Binning Algorithm to Recover Genomes from Multiple Metagenomic Datasets.” Bioinformatics 32 (4): 605–7. 10.1093/bioinformatics/btv638.

111. Xie, Li, and Wenying Shou. 2021. “Steering Ecological-Evolutionary Dynamics to Improve Artificial Selection of Microbial Communities.” Nature Communications 12 (1): 6799. 10.1038/s41467-021-26647-4.

112. Xu, Ling, and Devin Coleman-Derr. 2019. “Causes and Consequences of a Conserved Bacterial Root Microbiome Response to Drought Stress.” Current Opinion in Microbiology 49 (June): 1–6. 10.1016/j.mib.2019.07.003.

113. Xu, Ling, Zhaobin Dong, Dawn Chiniquy, Grady Pierroz, Siwen Deng, Cheng Gao, Spencer Diamond, et al. 2021. “Genome-Resolved Metagenomics Reveals Role of Iron Metabolism in Drought-Induced Rhizosphere Microbiome Dynamics.” Nature Communications 12 (1): 3209. 10.1038/s41467-021-23553-7.

114. Xu, Ling, Dan Naylor, Zhaobin Dong, Tuesday Simmons, Grady Pierroz, Kim K. Hixson, Young-Mo Kim, et al. 2018. “Drought Delays Development of the Sorghum Root Microbiome and Enriches for Monoderm Bacteria.” Proceedings of the National Academy of Sciences of the United States of America 115 (18): E4284–93. 10.1073/pnas.1717308115.

115. Xu, Peng, and Ertao Wang. 2023. “Diversity and Regulation of Symbiotic Nitrogen Fixation in Plants.” Current Biology: CB 33 (11): R543–59. 10.1016/j.cub.2023.04.053.

116. Yandigeri, Mahesh S., Kamlesh Kumar Meena, Divya Singh, Nityanand Malviya, Dhananjaya P. Singh, Manoj Kumar Solanki, Arvind Kumar Yadav, and Dilip K. Arora. 2012. “Drought-Tolerant Endophytic Actinobacteria Promote Growth of Wheat (Triticum Aestivum) under Water Stress Conditions.” Plant Growth Regulation 68 (3): 411–20. 10.1007/s10725-012-9730-2.

117. Zhang, Jingying, Weidong Liu, Jingshu Bu, Yanbing Lin, and Yang Bai. 2023. “Host Genetics Regulate the Plant Microbiome.” Current Opinion in Microbiology 72 (April): 102268. 10.1016/j.mib.2023.102268.

118. Zhuang, Lubo, Yan Li, Zhenshuo Wang, Yue Yu, Nan Zhang, Chang Yang, Qingchao Zeng, and Qi Wang. 2021. “Synthetic Community with Six Pseudomonas Strains Screened from Garlic Rhizosphere Microbiome Promotes Plant Growth.” Microbial Biotechnology 14 (2): 488–502. 10.1111/1751-7915.13640.

